# The Impact of Tokenizer Selection in Genomic Language Models

**DOI:** 10.1101/2024.09.09.612081

**Authors:** LeAnn M. Lindsey, Nicole L. Pershing, Anisa Habib, Keith Dufault-Thompson, W. Zac Stephens, Anne J. Blaschke, Xiaofang Jiang, Hari Sundar

## Abstract

Genomic language models have recently emerged as a new method to decode, interpret, and generate genetic sequences. Existing genomic language models have utilized various tokenization methods, including character tokenization, overlapping and non-overlapping k-mer tokenization, and byte-pair encoding, a method widely used in natural language models. Genomic sequences differ from natural language because of their low character variability, complex and overlapping features, and inconsistent directionality. These features make sub-word tokenization in genomic language models significantly different from both traditional language models and protein language models. This study explores the impact of tokenization in genomic language models by evaluating their downstream performance on forty-four classification fine-tuning tasks. We also perform a direct comparison of byte pair encoding and character tokenization in Mamba, a state-space model. Our results indicate that character tokenization outperforms sub-word tokenization methods on tasks that rely on nucleotide level resolution, such as splice site prediction and promoter detection. While byte-pair tokenization had stronger performance on the SARS-CoV-2 variant classification task, we observed limited statistically significant differences between tokenization methods on the remaining downstream tasks.

## Introduction

Tokenization is a fundamental step in the language model preprocessing pipeline and is used to parse an input sequence into segments called *tokens* that represent either words, subwords, or characters. These tokens are assigned numeric values and used as inputs to a neural network in order to learn context-specific embeddings. While sub-word tokenization methods have become a de facto standard in large language models (1, 2), various genomic language models (gLMs) have adopted different tokenization approaches, with no consensus emerging in the field. As genomic language models continue to advance, it will be essential to understand the performance impact of tokenization on downstream biological tasks.

Current genomic language models have been trained using three main tokenization methods: *character based tokenization*, in which the sequences are tokenized by individual nucleotides, *k-mer tokenization*, where the input is tokenized into either overlapping or non-overlapping substrings of length *k*, and sub-word tokenization using *byte-pair encoding* (Figure 1).

**Fig. 1.**
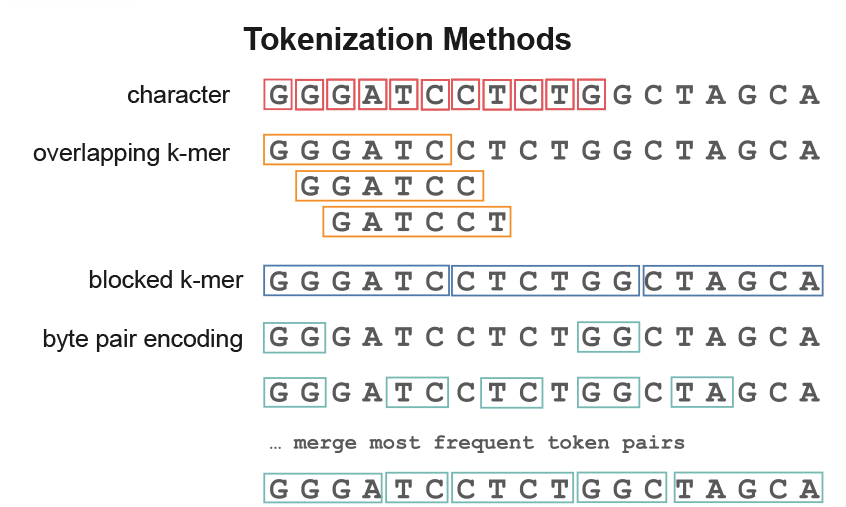
Tokenization of a sample input with character, overlapping and non-overlapping k-mer tokenization and byte-pair encoding.

Byte pair encoding (BPE) was originally developed as a method of compression and was later widely adopted by the natural language processing (NLP) community as a method of tokenization for language models. BPE iteratively calculates the frequency of adjacent characters and merges the most frequent pairs, using these frequencies to construct a vocabulary of the most frequently seen sub-words in a training corpus. This creates a compact representation that allows models to capture semantic relationships, reduce vocabulary size, and handle rare and out-of-vocabulary words. BPE also provides significant text compression benefits, with the DNABERT-2 tokenizer achieving 4-5x compression ratios (3). The non-overlapping k-mer tokenization used by the Nucleotide Transformer model, which we label *blocked k-mer*, also provides this same compression advantage. This efficient representation expands the effective context window capacity of the model, reducing training time and, subsequently, the cost to train the model.

Though subword tokenization provides many benefits, it also introduces several challenges that have been shown to adversely affect model performance in natural language models. Inconsistent tokenization, where the same word can be tokenized differently based on its position in the text, can lead to performance degradation, model hallucinations (4, 5), and a loss of semantic relationships between subwords (6). Lack of character-level transparency when using subword tokenization can also cause difficulty with tasks that require character-level reasoning such as precise spelling, letter counting, and arithmetic (5, 7).

Despite sub-word tokenization being broadly used in natural language processing, the ideal tokenization method is still being actively explored, with substantial research dedicated to understanding the impact of various tokenization choices (4, 8–11). Similar studies are needed in the genomic language domain, where the content and complexity differ significantly from natural language. Compared to natural language and protein sequences, nucleotide sequences have low character variability, contain overlapping and nested regulatory features, and have no obvious “word” demarcations. These differences suggest that the commonly used approaches for tokenization in natural language need to be evaluated in the significantly different context of nucleotide sequences.

The most significant work studying tokenization in biological language models to date is Dotan et al. (12), which tested five different tokenizers of various sizes: BPE (13), Unigram (14), WordPiece (15), characters, and pairs, against eight different biological datasets. They concluded that the choice of tokenizer has a significant impact on the downstream accuracy of the model, observing a change in accuracy as much as 5% and a change in MCC score as large as 0.10 depending on the tokenizer/task combination. Their experiments primarily focused on models trained on amino acid sequences, with only one experiment using nucleotide sequences as input, leaving open questions about the downstream impact of tokenization on nucleotide sequences.

Various researchers have discussed the impact of tokenization when introducing new gLMs, providing anecdotal evidence about the benefits and drawbacks of different approaches. The authors of DNABERT-2 compared BPE and k-mer tokenization in an ablation study, concluding that BPE performed better on average, although their experiments did indicate that k-mer tokenization had better performance in some tasks, including promoter detection (3). The authors of HyenaDNA (16) compared BPE with k-mer tokenization and concluded that using BPE tokenization with their model degraded their results. Schiff et al. (17) observed that with k-mer tokenization, small changes in the input sequence can result in dramatic changes in tokenization, but published no experiments linking these tokenization changes to downstream performance. (17). These studies demonstrate that tokenization choices can have significant downstream impact on model performance, highlighting the need for a systematic analysis of tokenization in the genomic language domain.

In this study, we investigate the following questions:

- Is sub-word tokenization in gLMs simply a form of compression, or does tokenization assist the model in capturing meaningful contextual relationships between tokens?
- How does the choice of tokenizer impact model performance on downstream tasks?

In order to investigate these questions, we compared three attention-based gLMs, three state space gLMs, and two baseline models on forty-four downstream classification tasks compiled from three published genomic benchmarks, the Genomic Benchmark (GB) (18), the Nucleotide Transformer Tasks (revised) (NTTv2) (19) and the Genome Understanding Evaluation (GUE) (3). We also trained a 4-layer Mamba state space model using both byte-pair encoding and character tokenization to gauge the direct impact of tokenization choice on model performance.

The results of our study indicate that the choice of tokenization method can significantly impact model performance, particularly on tasks that require the detection of specific nucleotide motifs. This work highlights the need for tailored machine learning approaches for biological applications that optimize pairing between the best suited tokenization methods and the biological feature(s) being investigated.

## Materials & Methods

### Genomic Language Models

We tested two baseline models, a simple three-layer CNN (18) trained using one-hot encoding and GPT-Neo-125m from EleutherAI. Although the GPT-Neo model was not trained on DNA, it performed significantly better than random so we include it as a second baseline comparison. GPT-Neo uses BPE tokenization, and no changes were made to the model or tokenizer to adapt them to DNA for these experiments.

We compared the transformer-based models, Nucleotide Transformer version 2 (nucleotide-transformer-v2-500mmulti-species) (19), DNABERT (20), and DNABERT-2 (3), against three of the state space genomic language models, HyenaDNA (16), Mamba (21), and Caduceus (17). To directly compare the effect of a tokenizer on a state space model, we pretrained a four layer Mamba model using an input sequence length of 4096, dimension 256, using both character and BPE tokenization. More detail on this model and our hyperparameter tuning during pretraining is available in the Supplementary Material. All of the state space models were trained using the same human reference genome (Hg38) dataset used to train the Caduceus model (17) and used a context length of 4000 nucleotides.

### Benchmarks

We benchmarked all models against three published genomic fine-tuning benchmarks: the recently revised Nucleotide Transformer Tasks (19), the Genomic Benchmark (18), and the Genome Understanding Evaluation (3). All fine-tuning tasks were replicated a minimum of ten times. A hyperparameter search for the best batch size and learning rate was performed using three initial seed values for the state space models since they are sensitive to these parameters. For the attention-based models, we used the learning rate and batch sizes reported in the fine tuning experiments in Zhou et al (22). Details are available in the Supplementary Material.

### Metrics

The mean accuracy and Mathews Correlation Coefficient (MCC) were reported on each fine tuning task.

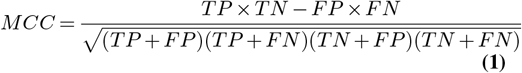

The MCC score ranges from -1 to 1 with 1 representing perfect prediction and 0 representing random performance. In the case of tasks that have more than two labels, the macro average was reported, which gives equal weight to each class regardless of the number of members of each class.

### Statistical Testing

To evaluate the significance of the performance differences between the Mamba-char and Mamba-bpe models, we performed paired t-tests, pairing replicates with identical random seeds. We performed tests at both the task and task category aggregation levels. To address multiple testing concerns arising from categorical comparisons, we applied Bonferroni correction (23) at each level to control the family-wise error rate. For individual task comparisons we used *α* = 0.05*/*number of tasks and for category-level comparisons, we used *α* = 0.05*/*number of categories. Bonferroni correction is a conservative approach that reduces the significance threshold to prevent inflation of statistical significance that can occur when multiple tests are performed on the same dataset.

### Tokenizers

Our experiments on tokenization were implemented using the transformers library from *HuggingFace* (24). We limited this study to token vocabularies of size 4096 because of the experiments by Zhou et al. (3) that recommended tokenized vocabularies of this size for genomic language models.

### Genomic Tasks

The genomic tasks in the benchmarking datasets can be grouped into nine categories: regulatory elements, promoter detection, enhancer detection, transcription factor binding site prediction, epigenetic marks prediction, splice site detection, coding region detection, taxonomic classification and virus variant detection.

The regulatory elements category includes Ensembl reference datasets annotating promoters, enhancers, and open chromatin regions (25, 26). Promoters are regions typically located upstream of gene transcription start sites and serve as platforms for RNA polymerase and transcription factors to initiate gene expression; these can be further sub-classified by specific genomic sequence features. Enhancers are regulatory elements that can be located upstream, downstream, or within the introns of their target genes, often acting over long distances. Open chromatin regions are genomic regions accessible for protein binding that lack evidence to classify them as promoters or enhancers.

Transcription factor binding sites contain motifs ranging from 6 to 20 nt in length, where transcription factors bind to regulate gene expression. They are usually located within promoter, enhancer, or other regulatory regions. These motifs typically display sequence flexibility, allowing inexact matches to the consensus motif, with the binding affinity correlating with the similarity to the optimal binding motif. (27) Open chromatin regions are genomic regions that are accessable to DNA-binding proteins due to relaxed nucleosome packing. Chromatin accessibility and nucleosome packing are regulated by multiple epigenetic mechanisms, including post-translational modification of histone tails (such as acetylation and methylation), which alter nucleosome structure and chromatin accessibility. The epigenetic marks prediction task focuses on identifying regions associated with specific histone modifications that regulate expression.

Splice sites are specific sequences at exon-intron boundaries that direct the splicing machinery and typically include canonical GT dinucleotides at the 5’ donor sites and AG dinucleotides at the 3’ acceptor sites.

Tasks that did not align with other genomic feature categories –including coding vs intergenic, worm vs human and SARSCoV-2 variant classification– were maintained as standalone categories.

## Results

### Tokenization Differentially Impacts Performance of gLMs on Specific Benchmark Tasks

The best performing model in each task category varied significantly, with the Nucleotide Transformer having the highest performance on the splice sites and epigenetic marks detection tasks (Figure 2, Table 2). The original DNABERT model had the highest MCC score on the promoter detection task, Caduceus had the highest performance on regulatory elements, the enhancer detection and transcription factor binding site prediction tasks. The Mamba-bpe model had the best performance on the SARS-CoV-2 virus variant detection task. Caduceus had the best performance overall, followed closely by DNABERT2.

In the direct comparison experiment using the Mamba model, the different tokenization approaches showed significantly different performance on a subset of the biological tasks (Figure 3, Tables 3, 4.) In the promoters category, the mean difference was 0.0289 (*t* = 6.09, *p <* 0.0001) with character tokenization having better performance on this task. In the splice site detection category, character tokenization shows significantly better performance with a mean MCC difference of 0.2235 (*t* = 12.44, *p <* 0.0001). In the virus variant detection task, byte-pair encoding had better performance with a mean difference of 0.0726 (*t* = 7.17, *p <* 0.0002). We observe a slight but statistically significant difference in favor of character tokenization on the epigenetic marks and taxonomic task categories. In the coding, enhancers and transcription factor categories, we do not observe a statistically significant difference between tokenizers.

**Table 1.**
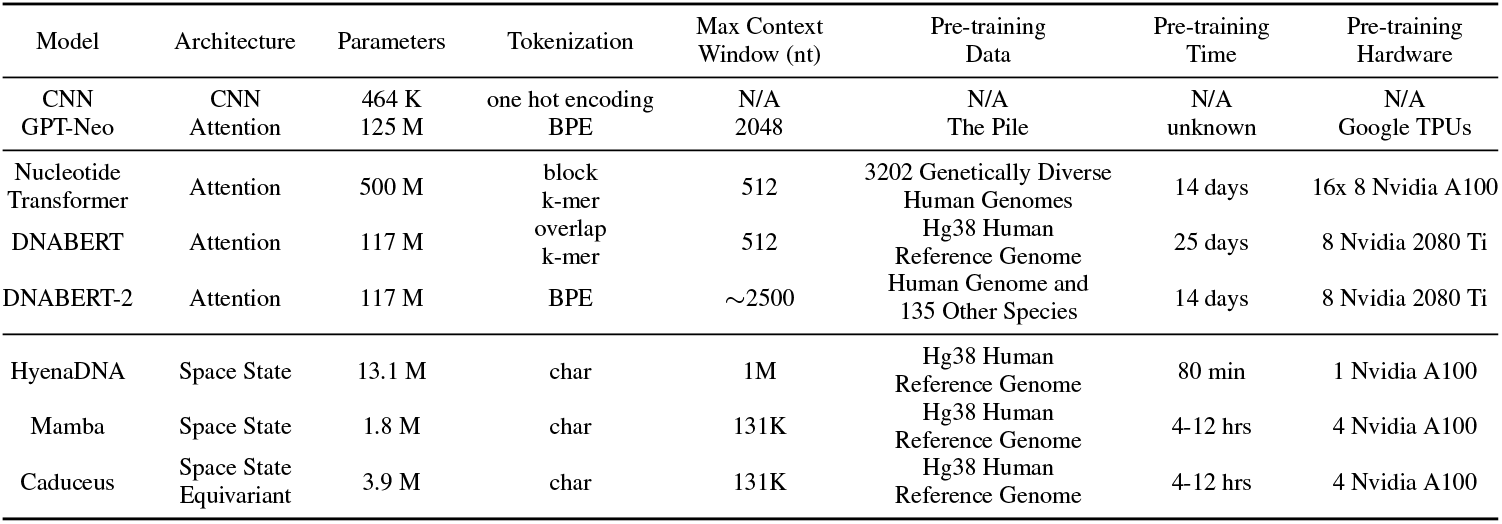
Details of all models used in the benchmarking experiments. A simple 3-layer CNN was used as a baseline model. The GPT-Neo pre-trained model was used as a second baseline. The Time and Hardware columns specify the hardware used and total time needed for pretraining.

**Table 2.** Overview of MCC scores across models summarized by task category and benchmark. The highest performing model in each row is highlighted in bold and underlined. The benchmarks included are: Genomic Benchmark (GB) (18), Nucleotide Transformer Tasks (NTT) (19), GUE (Genome Understanding Evaluation) (3)

| Category | CNN | GPT-2 | DNABERT-2 (bpe) | Mamba (bpe) | NT (blocked k-mer) | DNABERT (k-mer) | HyenaDNA (char) | Mamba (char) | Caduceus (char) |
| --- | --- | --- | --- | --- | --- | --- | --- | --- | --- |
| model size (parameters) | 464K | 125M | 117M | 3.9M | 500M | 117M | 13.1M | 1.8M | 3.9M |
| regulatory | 0.638 | 0.568 | 0.664 | 0.760 | 0.634 | 0.431 | 0.760 | 0.716 | **<u>0.778</u>** |
| promoters | 0.749 | 0.716 | 0.762 | 0.639 | 0.754 | **<u>0.777</u>** | 0.675 | 0.667 | 0.728 |
| enhancers | 0.467 | 0.457 | 0.559 | 0.539 | 0.520 | 0.441 | 0.552 | 0.548 | **<u>0.573</u>** |
| transcription factors | 0.496 | 0.557 | 0.685 | 0.616 | 0.490 | 0.635 | 0.667 | 0.605 | **<u>0.703</u>** |
| epigenetic marks | 0.480 | 0.504 | 0.568 | 0.460 | **<u>0.576</u>** | 0.526 | 0.481 | 0.470 | 0.501 |
| splice sites | 0.893 | 0.738 | 0.835 | 0.631 | **<u>0.941</u>** | 0.931 | 0.884 | 0.854 | 0.870 |
| virus variant detection | 0.128 | 0.549 | 0.401 | **<u>0.652</u>** | 0.498 | 0.389 | 0.620 | 0.576 | 0.630 |
| GUE | 0.569 | 0.614 | 0.699 | 0.618 | 0.587 | 0.682 | 0.671 | 0.621 | **<u>0.707</u>** |
| Genomic Benchmark | 0.622 | 0.570 | 0.702 | 0.719 | 0.625 | 0.502 | 0.729 | 0.705 | **<u>0.750</u>** |
| Nucleotide Transformer Tasks | 0.606 | 0.579 | 0.643 | 0.527 | **<u>0.684</u>** | 0.638 | 0.588 | 0.585 | 0.608 |
| Overall | 0.594 | 0.583 | 0.673 | 0.599 | 0.634 | 0.617 | 0.643 | 0.621 | **<u>0.674</u>** |
|  | Baseline |  | Sub-word Tokenization |  |  | Nucleotide Level Tokenization |  |  |  |

**Table 3.**
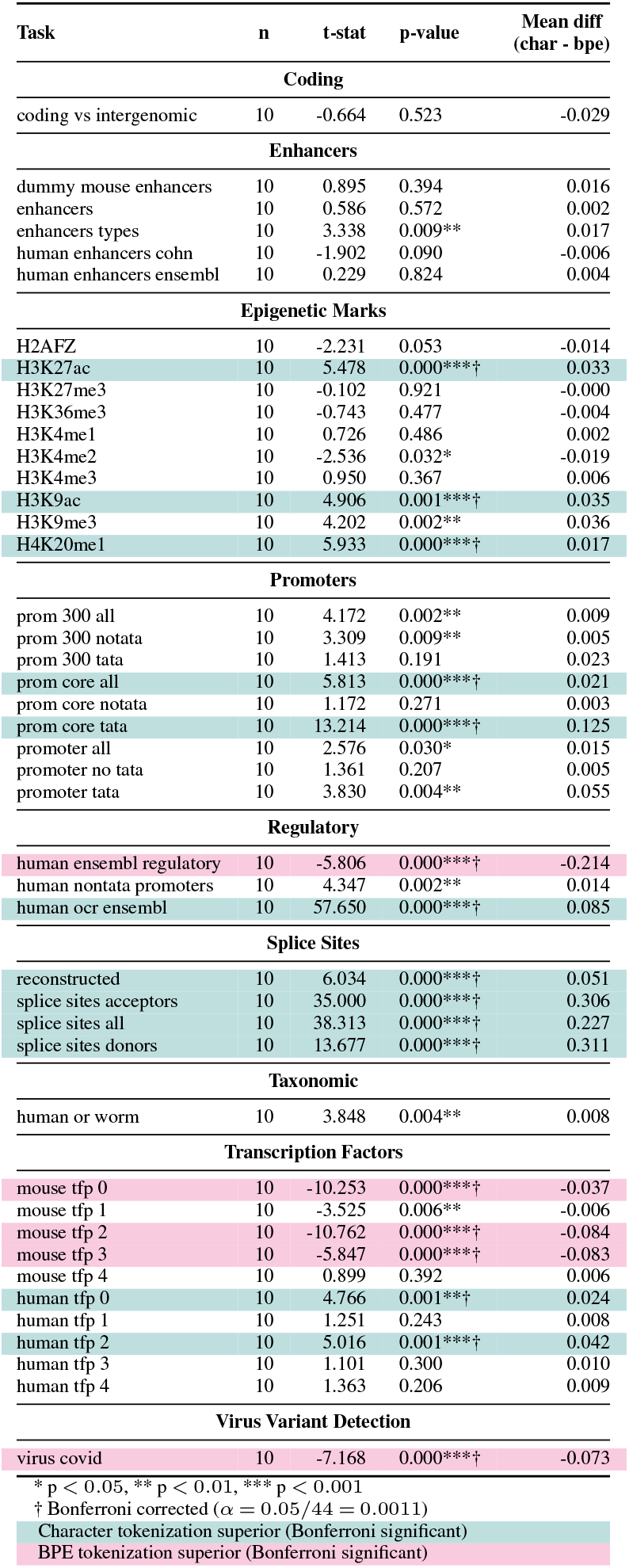
Task-Level Paired Comparison of Character-Level vs. Byte Pair Encoding Tokenization on MCC Scores in a 4 layer Mamba-DNA Model (Matched by Seed)

**Table 4.**
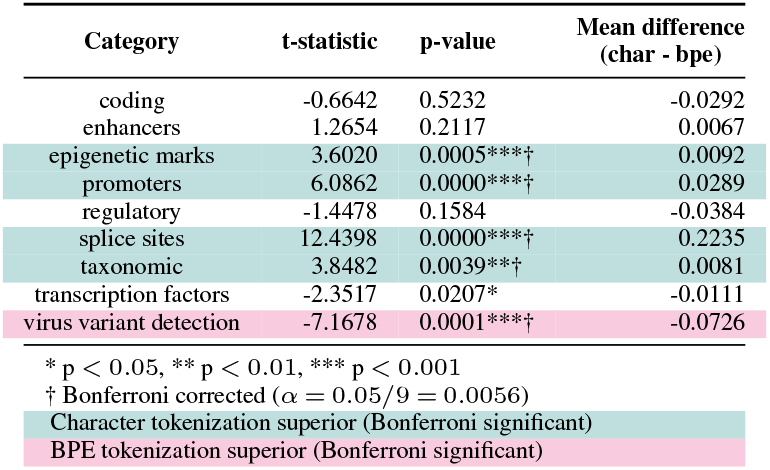
Paired Comparison of Character-Level vs. Byte Pair Encoding Tokenization on MCC Scores Across Different Genomic Features in a 4 layer Mamba-DNA Model (Matched by Seed)

**Fig. 2.**
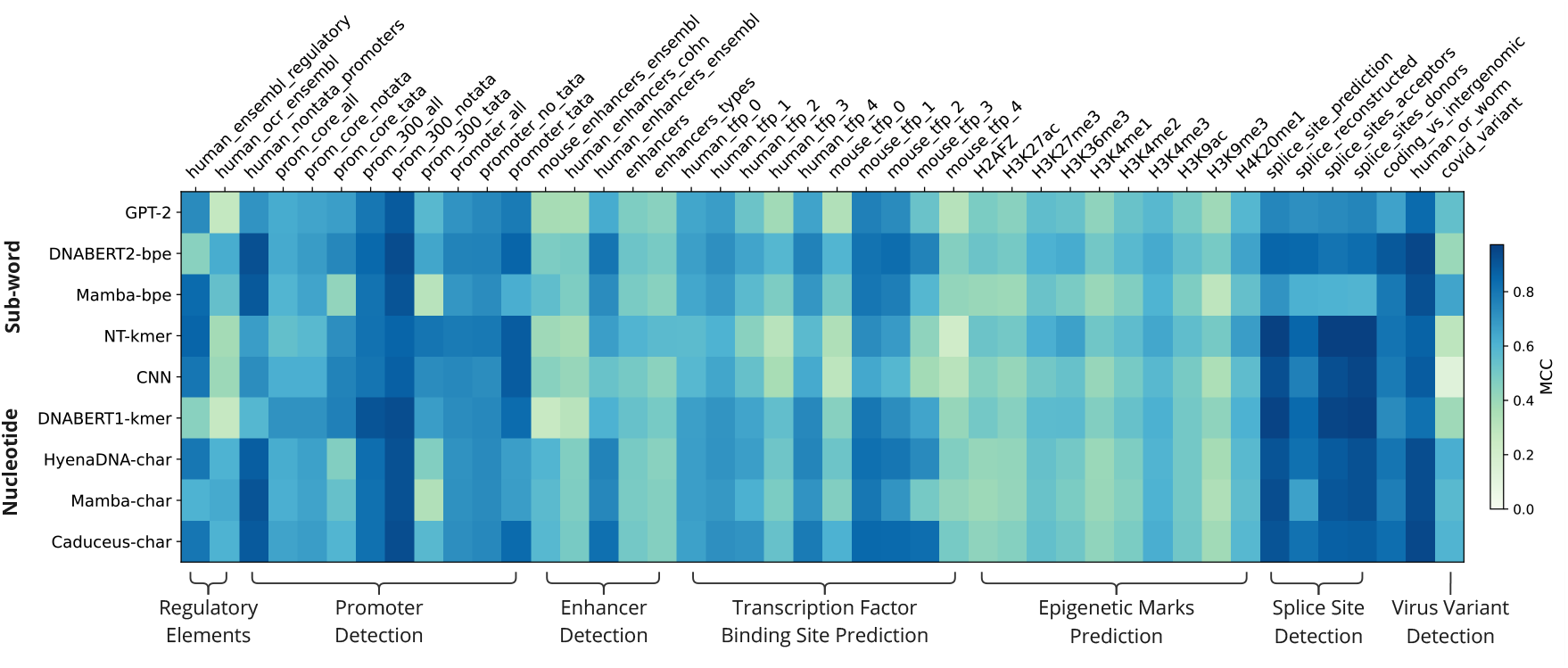
Model Performance as measured by the Matthews Correlation Cooefficient on all Benchmark Tasks, grouped by task category. MCC score is visualized as a blue-green color gradient, with darker blue indicating better performance. The rows are clustered by the type of tokenization method, with sub-word tokenization near the top followed by the nucleotide level tokenization methods.

**Fig. 3.**
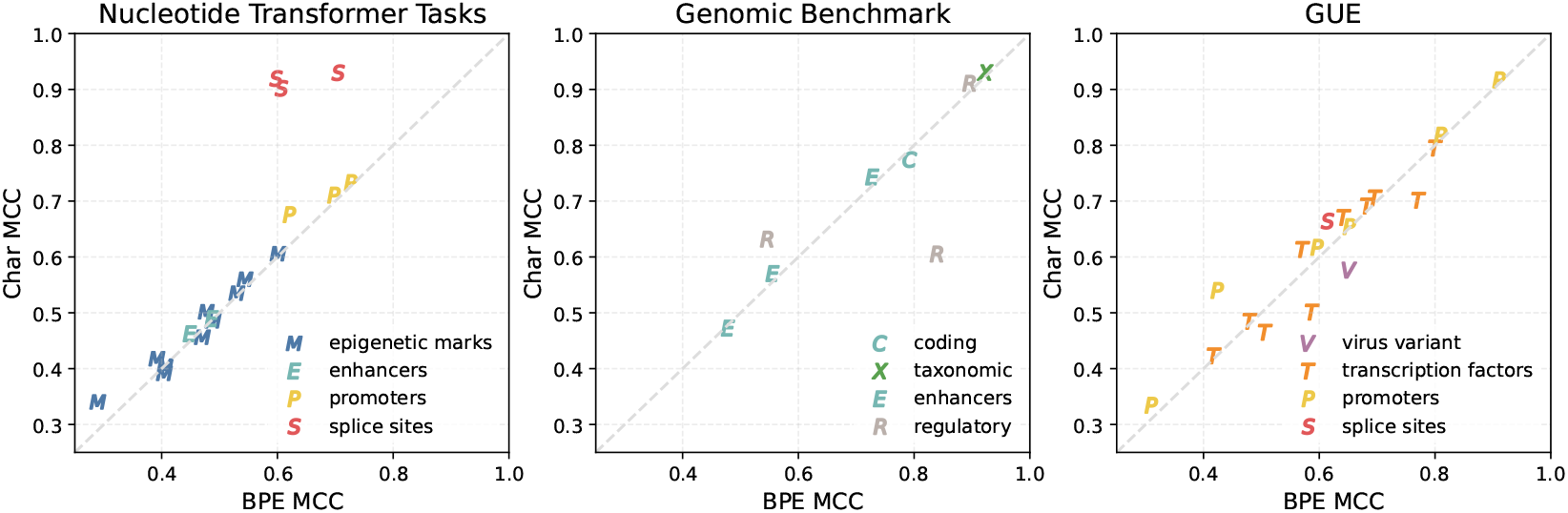
Comparison of tokenization methods in a four layer Mamba-DNA model on all three genomic benchmarks. Markers above the diagonal line indicate that the Mamba model with character tokenization has higher MCC score than BPE tokenization on the same dataset. Markers are colored by task category.

### Limited Overlap Between Learned BPE Tokens and Known Regulatory Motifs

In order to determine if the vocabulary learned by the byte-pair encoding process contains known regulatory motifs, we did a comparison with exact string matching between the BPE tokenizer vocabulary and the JASPAR 2024 CORE transcription factor binding motif database (28) and found that only 1.54% of the learned tokens correspond to annotated regulatory motifs.

## Discussion

While tokenization approaches have been extensively investigated and benchmarked in natural language models, their impacts on genomic language models have remained largely unexplored. The distinct characteristics of genomic language relative to natural language necessitate a careful evaluation of the advantages and limitations of various tokenization strategies when applied to genomic data. Our comparison of tokenization approaches on a range of biological tasks provide insights into the importance of modeling choices in machine learning and can guide future model development in the biological sciences.

Our direct comparison of tokenization methods using the Mamba architecture indicates that single-nucleotide resolution is superior for specific downstream tasks, notably promoter detection and splice site prediction. This aligns with research in the NLP community that demonstrates that character-level resolution improves performance on downstream tasks such as spelling and arithmetic tasks (7, 11, 29, 30). In the case of splice site prediction, canonical splice sites have highly conserved dinucleotide sequences (e.g., GT/AG) at exon-intron boundaries, and a single nucleotide mutation in this region can disrupt splicing function (31). Similarly, promoters often contain specific nucleotide motifs which can be highly conserved within and between species (32) and have conserved spatial arrangements in the nucleotide sequences (33). The mean MCC differences that we see between the tokenization methods for promoter detection (Δ = +0.0289) and particularly for splice-site detection (Δ = +0.2235) suggest that at this relatively small parameter size, without the single nucleotide precision of character tokenization, the model’s ability to predict these features is reduced.

The attention-based models that used sub-word tokenization perform better on the splice site and promoter detection tasks than the Mamba-BPE model, which may indicate that the attention mechanism better compensates for sub-word tokenization’s reduced nucleotide-level precision when identifying specific sequence motifs.

The original DNABERT model scored 0.931 on the splice site detection task, very close to the highest performing model, Nucleotide Transformer (0.941), which has significantly more parameters and is trained on a more diverse dataset. Although the overlapping k-mer tokenization used by DNABERT is not character-level tokenization, the single-nucleotide shift between adjacent tokens preserves nucleotide-level information content, making it effectively equivalent to character-level tokenization, and allowing the model to learn at a nucleotide level resolution. The DNABERT-2 model, which uses BPE tokenization, drops in performance on the same category by 0.096, suggesting that nucleotide-level resolution improves performance in splice site prediction. We postulate that the larger parameter size of the Nucleotide Transformer enables the model to learn more splice site patterns, and its consistent token size (6 nt) may also facilitate the model’s learning of specific distances.

In the Mamba model direct comparison, byte-pair encoding outperforms character tokenization on the SARS-CoV-2 variant classification task. This task challenges the model to differentiate between nine different variant classes, while all other classification tasks have only two or three classes. This could indicate that there are other categories of tasks not tested in this study where BPE may provide significant advantage.

The regulatory and transcription factor tasks had mixed results, with some datasets favoring BPE and some datasets favoring character tokenization, but there was no statistically significant difference at the category level.

In all remaining task categories, we did not observe any meaningful differences in performance. We hypothesize that these tasks likely rely on broader trends in sequence composition. Sequences from different organisms, for example, are not primarily differentiated by specific nucleotide motifs. Instead, other factors like GC content and oligonucleotide frequency distributions have been found to be significantly different between organisms (34). Similarly, prediction of the sites of epigenetic modification of histones has been associated with broader trends in sequence composition rather than the presence of specific motifs (35). The comparable performance of different tokenizers on these tasks suggests that when predictions depend on broad compositional patterns rather than specific motifs, multiple tokenization approaches can effectively capture the relevant features.

Our motif analysis indicated that the byte-pair encoding algorithm, as it is currently being applied to genomic sequences, does not efficiently learn a vocabulary of known genomic motifs. One probable explanation is that although motifs are important, they are not frequent, and the BPE algorithm builds the vocabulary based on the most frequent words that appear in the training dataset. The training sets used for the published gLMs are primarily made up of randomly selected DNA segments. Curating these training sets to contain a higher proportion of known regulatory motifs or regions may improve the model’s ability to learn these motifs.

### Limitations and Future Work

To limit the scope of this study we did not investigate different vocabulary sizes for the sub-word tokenizers, and did not explore every available subword tokenizer. We focused on the tokenizers currently used in published pre-trained models and fixed the vocabulary size at 4096 based on the recommendation of Zhou et al. (3). We tested only classification tasks and no regression or generation tasks were compared. The short context window in current attention-based genomic language models precluded us from a direct comparison between character based and byte pair encoding, however, with new model architectures expanding this context window, a direct comparison of these tokenization methods in an attention-based model should be completed. In addition, compared to large language models, the model sizes of all published genomic language models are relatively small, and they have been trained primarily on eukaryotic species. More study is needed to evaluate how tokenization decisions will affect models with significantly more parameters and models trained on data from other taxonomic domains. The focus of this study was tokenization, but our results illustrate significant performance differences between model architecture decisions, including differences between the different state space model architectures. These differences should be explored more fully in future work.

### Conclusion

In conclusion, our experiments demonstrate that the selection of tokenization methods substantially influences model performance on downstream genomic tasks. The performance of the BPE tokenizer on the difficult nine category discriminatory task of SARS-CoV-2 variant classification illustrates that on more challenging genomic tasks, BPE tokenizers may have an advantage beyond compression.

We acknowledge that the limited performance of BPE in our study could also be attributed to the limited parameter size of current genomic language models. Although BPE has proven valuable in natural language processing, our results suggest that it may not be optimal for all genomic classification tasks, particularly those that require the precise identification of biological motifs. This work underscores the critical role of domain-specific knowledge in model development and highlights the necessity for further investigation into genomic language model tokenization, challenging assumptions carried over from natural language processing. Ultimately, these findings emphasize the potential to develop novel tokenization strategies tailored to the unique characteristics of genomic sequences, potentially incorporating biological priors or adaptive schemes that preserve biologically relevant units, to achieve improved performance.

## Acknowledgments

This work utilized the computational resources of the NIH HPC Biowulf cluster (https://hpc.nih.gov), computational resources and support from the Center for High Performance Computing at the University of Utah as well as Bridges-2 at Pittsburgh Supercomputing Center and Delta at the National Center for Supercomputing Applications (NCSA) through allocation BIO230092 from the Advanced Cyberinfrastructure Coordination Ecosystem: Services & Support (ACCESS) program, which is supported by National Science Foundation grants #2138259, #2138286, #2138307, #2137603, and #2138296. L.L, K.D. and X.J. are supported by the Division of Intramural Research (DIR) of the National Library of Medicine (NLM), National Institutes of Health. L.L, A.H. and H.S. are supported by funds from the National Science Foundation (NSF: #2222322). N.P. was supported by the National Center for Advancing Translational Sciences of the National Institutes of Health under Award Numbers UM1TR004409 and 1K12TR004413. The content is solely the responsibility of the authors and does not necessarily represent the official views of the National Institutes of Health. We would like also to acknowledge and thank Yair Schiff for his technical support for the state space models and Zhihan Zhou for his technical support for DNABERT-2.

## References

1. Martin Berglund and Brink van der Merwe. Formalizing BPE tokenization. Workshop on Non-Classical Models for Automata and Applications, 388:16–27, September 2023. doi: 10.4204/eptcs.388.4.

2. Anh Khoa Ngo Ho and François Yvon. Optimizing word alignments with better subword tokenization. In The 18th biennial conference of the International Association of Machine Translation, Proceedings of the 18th Biennial Machine Translation Summit :Volume 1: Research Track, Miami (virtual), United States, August 2021.

3. Zhihan Zhou, Yanrong Ji, Weijian Li, Pratik Dutta, Ramana Davuluri, and Han Liu. DNABERT-2: efficient foundation model and benchmark for multi-species genome, June 2023. arXiv: 2306.15006.

4. Kaiser Sun, Qi Pan, Yuhao Zhang, Lan Liu, William Yang Wang, and Zhiheng Huang. Tokenization consistency matters for generative models on extractive NLP tasks. Conference on Empirical Methods in Natural Language Processing, December 2022. doi: 10.48550/arxiv.2212.09912. 10.48550/arxiv.2212.09912.

5. Kaj Bostrom and Greg Durrett. Byte Pair Encoding is Suboptimal for Language Model Pretraining. In Trevor Cohn, Yulan He, and Yang Liu, editors, Findings of the Association for Computational Linguistics: EMNLP 2020, pages 4617–4624, Online, November 2020. Association for Computational Linguistics. doi: 10.18653/v1/2020.findings-emnlp.414.

6. Khuyagbaatar Batsuren, Ekaterina Vylomova, Verna Dankers, Tsetsuukhei Delgerbaatar, Omri Uzan, Yuval Pinter, and Gábor Bella. Evaluating Subword Tokenization: Alien Subword Composition and OOV Generalization Challenge, April 2024. 2404.13292 [cs].

7. Yekun Chai, Yewei Fang, Qiwei Peng, and Xuhong Li. Tokenization Falling Short: On Subword Robustness in Large Language Models. In Yaser Al-Onaizan, Mohit Bansal, and Yun-Nung Chen, editors, Findings of the Association for Computational Linguistics: EMNLP 2024, pages 1582–1599, Miami, Florida, USA, November 2024. Association for Computational Linguistics. doi: 10.18653/v1/2024.findings-emnlp.86.

8. Craig W Schmidt, Varshini Reddy, Haoran Zhang, Alec Alameddine, Omri Uzan, Yuval Pinter, and Chris Tanner. Tokenization Is More Than Compression. In Yaser Al-Onaizan, Mohit Bansal, and Yun-Nung Chen, editors, Proceedings of the 2024 Conference on Empirical Methods in Natural Language Processing, pages 678–702, Miami, Florida, USA, November 2024. Association for Computational Linguistics. doi: 10.18653/v1/2024.emnlp-main.40.

9. Phillip Rust, Jonas Pfeiffer, Ivan Vulić, Sebastian Ruder, and Iryna Gurevych. How good is your tokenizer? On the monolingual performance of multilingual language models. In Chengqing Zong, Fei Xia, Wenjie Li, and Roberto Navigli, editors, Proceedings of the 59th Annual Meeting of the Association for Computational Linguistics and the 11th International Joint Conference on Natural Language Processing (Volume 1: Long Papers), pages 3118– 3135, Online, August 2021. Association for Computational Linguistics. doi: 10.18653/v1/2021.acl-long.243.

10. Nived Rajaraman, Jiantao Jiao, and Kannan Ramchandran. Toward a theory of tokenization in LLMs, April 2024. 2404.08335.

11. Aaditya K. Singh and D. J. Strouse. Tokenization counts: the impact of tokenization on arithmetic in frontier LLMs. arXiv preprint 2402.14903, 2024.

12. Edo Dotan, Gal Jaschek, Tal Pupko, and Yonatan Belinkov. Effect of tokenization on transformers for biological sequences. Bioinformatics, 40(4):btae196, April 2024. ISSN 1367-4811. doi: 10.1093/bioinformatics/btae196.

13. Rico Sennrich, Barry Haddow, and Alexandra Birch. Neural Machine Translation of Rare Words with Subword Units. In Katrin Erk and Noah A. Smith, editors, Proceedings of the 54th Annual Meeting of the Association for Computational Linguistics (Volume 1: Long Papers), pages 1715–1725, Berlin, Germany, August 2016. Association for Computational Linguistics. doi: 10.18653/v1/P16-1162.

14. Taku Kudo. Subword Regularization: Improving Neural Network Translation Models with Multiple Subword Candidates, 2018. _eprint: 1804.10959.

15. Mike Schuster and Kaisuke Nakajima. Japanese and Korean voice search. In 2012 IEEE International Conference on Acoustics, Speech and Signal Processing (ICASSP), pages 5149–5152, 2012. doi: 10.1109/ICASSP.2012.6289079.

16. Eric Nguyen, Michael Poli, Marjan Faizi, Armin W. Thomas, Callum Birch Sykes, Michael Wornow, Aman Patel, Clayton Rabideau, Stefano Massaroli, Yoshua Bengio, Stefano Ermon, Stephen A. Baccus, and Christopher Ré. HyenaDNA: long-range genomic sequence modeling at single nucleotide resolution. In Proceedings of the 37th International Conference on Neural Information Processing Systems, NIPS ‘23, Red Hook, NY, USA, 2023. Curran Associates Inc. event-place: New Orleans, LA, USA.

17. Yair Schiff, Chia-Hsiang Kao, Aaron Gokaslan, Tri Dao, Albert Gu, and Volodymyr Kuleshov. Caduceus: Bi-directional equivariant long-range DNA sequence modeling. arXiv preprint 2403.03234, 2024.

18. Katarína Grešová, Vlastimil Martinek, David Čechák, Petr Šimeček, and Panagiotis Alexiou. Genomic benchmarks: a collection of datasets for genomic sequence classification. BMC Genomic Data, 24(1):25, May 2023. ISSN 2730-6844. doi: 10.1186/s12863-023-01123-8.

19. Hugo Dalla-Torre, Liam Gonzalez, Javier Mendoza-Revilla, Nicolas Lopez Carranza, Adam Henryk Grzywaczewski, Francesco Oteri, Christian Dallago, Evan Trop, Bernardo P. de Almeida, Hassan Sirelkhatim, Guillaume Richard, Marcin Skwark, Karim Beguir, Marie Lopez, and Thomas Pierrot. The Nucleotide Transformer: building and evaluating robust foundation models for human genomics, October 2024. bioRxiv: 10.1101/2023.01.11.523679.

20. Yanrong Ji, Zhihan Zhou, Han Liu, and Ramana V Davuluri. DNABERT: pre-trained bidirectional encoder representations from transformers model for DNA-language in genome. Bioinformatics, 37:2112–2120, August 2021.

21. Albert Gu and Tri Dao. Mamba: Linear-Time Sequence Modeling with Selective State Spaces, May 2024. 2312.00752.

22. Zhihan Zhou, Yanrong Ji, Weijian Li, Pratik Dutta, Ramana Davuluri, and Han Liu. DNABERT-2: Efficient Foundation Model and Benchmark For Multi-Species Genome, March 2024. 2306.15006 [q-bio].

23. Olive Jean Dunn. Estimation of the Medians for Dependent Variables. The Annals of Mathematical Statistics, 30(1):192 – 197, 1959. doi: 10.1214/aoms/1177706374. Publisher: Institute of Mathematical Statistics.

24. Thomas Wolf, Lysandre Debut, Victor Sanh, Julien Chaumond, Clement Delangue, Anthony Moi, Pierric Cistac, Tim Rault, Rémi Louf, Morgan Funtowicz, Joe Davison, Sam Shleifer, Patrick von Platen, Clara Ma, Yacine Jernite, Julien Plu, Canwen Xu, Teven Le Scao, Sylvain Gugger, Mariama Drame, Quentin Lhoest, and Alexander M. Rush. HuggingFace’s Transformers: State-of-the-art natural language processing, July 2020. 1910.03771.

25. Kevin L Howe, Premanand Achuthan, James Allen, Jamie Allen, Jorge Alvarez-Jarreta, M Ridwan Amode, Irina M Armean, Andrey G Azov, Ruth Bennett, Jyothish Bhai, Konstantinos Billis, Sanjay Boddu, Mehrnaz Charkhchi, Carla Cummins, Luca Da Rin Fioretto, Claire Davidson, Kamalkumar Dodiya, Bilal El Houdaigui, Reham Fatima, Astrid Gall, Carlos Garcia Giron, Tiago Grego, Cristina Guijarro-Clarke, Leanne Haggerty, Anmol Hemrom, Thibaut Hourlier, Osagie G Izuogu, Thomas Juettemann, Vinay Kaikala, Mike Kay, Ilias Lavidas, Tuan Le, Diana Lemos, Jose Gonzalez Martinez, José Carlos Marugán, Thomas Maurel, Aoife C McMahon, Shamika Mohanan, Benjamin Moore, Matthieu Muffato, Denye N Oheh, Dimitrios Paraschas, Anne Parker, Andrew Parton, Irina Prosovetskaia, Manoj P Sakthivel, Ahamed I Abdul Salam, Bianca M Schmitt, Helen Schuilenburg, Dan Sheppard, Emily Steed, Michal Szpak, Marek Szuba, Kieron Taylor, Anja Thormann, Glen Threadgold, Brandon Walts, Andrea Winterbottom, Marc Chakiachvili, Ameya Chaubal, Nishadi De Silva, Bethany Flint, Adam Frankish, Sarah E Hunt, Garth R IIsley, Nick Langridge, Jane E Loveland, Fergal J Martin, Jonathan M Mudge, Joanella Morales, Emily Perry, Magali Ruffier, John Tate, David Thybert, Stephen J Trevanion, Fiona Cunningham, Andrew D Yates, Daniel R Zerbino, and Paul Flicek. Ensembl 2021. Nucleic Acids Research, 49(D1): D884–D891, January 2021. ISSN 0305-1048. doi: 10.1093/nar/gkaa942.

26. Daniel R. Zerbino, Steven P. Wilder, Nathan Johnson, Thomas Juettemann, and Paul R. Flicek. The ensembl regulatory build. Genome Biology, 16(1):56, March 2015. ISSN 1474-760X. doi: 10.1186/s13059-015-0621-5.

27. François Spitz and Eileen E. M. Furlong. Transcription factors: from enhancer binding to developmental control. Nature Reviews Genetics, 13(9):613–626, September 2012. ISSN 1471-0064. doi: 10.1038/nrg3207. Publisher: Nature Publishing Group.

28. I. Rauluseviciute, R. Riudavets-Puig, R. Blanc-Mathieu, J.A. Castro-Mondragon, K. Ferenc, V. Kumar, R.B. Lemma, J. Lucas, J. Chèneby, D. Baranasic, A. Khan, O. Fornes, S. Gundersen, M. Johansen, E. Hovig, B. Lenhard, A. Sandelin, W.W. Wasserman, F. Parcy, and A. Mathelier. JASPAR 2024: 20th anniversary of the open-access database of transcription factor binding profiles. Nucleic Acids Research, 52(D1):D174–D182, January 2024. doi:10.1093/nar/gkad1059.

29. Linting Xue, Aditya Barua, Noah Constant, Rami Al-Rfou, Sharan Narang, Mihir Kale, Adam Roberts, and Colin Raffel. ByT5: Towards a token-free future with pre-trained byte-to-byte models, March 2022. 2105.13626.

30. Yi Tay, Vinh Q. Tran, Sebastian Ruder, Jai Gupta, Hyung Won Chung, Dara Bahri, Zhen Qin, Simon Baumgartner, Cong Yu, and Donald Metzler. Charformer: Fast Character Transformers via Gradient-based Subword Tokenization. International Conference on Learning Representations, 2021. ARXIV_ID: 2106.12672 S2ID:e79d1206292bc5e67ba19737d87d4b2ea4a37105.

31. Ruohan Wang, Zishuai Wang, Jianping Wang, and Shuaicheng Li. SpliceFinder: ab initio prediction of splice sites using convolutional neural network. BMC Bioinformatics, 20(23):652, December 2019. ISSN 1471-2105. doi: 10.1186/s12859-019-3306-3.

32. Chuhu Yang, Eugene Bolotin, Tao Jiang, Frances M. Sladek, and Ernest Martinez. Prevalence of the initiator over the TATA box in human and yeast genes and identification of DNA motifs enriched in human TATA-less core promoters. Gene, 389(1):52–65, March 2007. ISSN 0378-1119. doi: 10.1016/j.gene.2006.09.029.

33. Aditi Kanhere and Manju Bansal. Structural properties of promoters: similarities and differences between prokaryotes and eukaryotes. Nucleic Acids Research, 33(10):3165–3175, June 2005. ISSN 0305-1048. doi: 10.1093/nar/gki627.

34. Samuel Karlin and Jan Mrázek. Compositional differences within and between eukaryoticgenomes. Proceedings of the National Academy of Sciences of the United States of America, 94(19):10227–10232, September 1997. ISSN 0027-8424.

35. Tho Hoan Pham, Tu Bao Ho, Dang Hung Tran, and Kenji Satou. Prediction of Histone Modifications in DNA sequences. In 2007 IEEE 7th International Symposium on BioInformatics and BioEngineering, pages 959–966, October 2007. doi: 10.1109/BIBE.2007.4375674.

