## Supplementary Material for "The Impact of Tokenizer Selection in Genomic Language Models"

### Supplemental Material: The Impact of Tokenizer Selection in Genomic Language Models

#### Acrynms

DNA - Deoxyribonucleic acid
NLP - Natural Language Processing
LMs - Language Models
pLMs - Protein Language Models
gLMs - Genomic Language Models
BPE - Byte Pair Encoding
OCR - Open Chromatin Region
CNN - Convolutional Neural Network
RNN - Recurrent Neural Network

#### Model Training Details

The CNN Baseline model used was the model specified in [5], as implemented in the Caduceus Github repository. For the GPT Baseline model we used EleutherAI/gpt-neo-125m, which is a EleutherAI's replication of the GPT-3 architecture. For DNABERT, and DNABERT-2, the pre-trained models provided by the authors were used (zhihan1996/DNA_bert_6, zhihan1996/DNABERT-2-117M). The InstaDeepAI/nucleotide-transformer-v2-500m-multi-species version of the Nucleotide Transformer was chosen as it was the largest multi-species model that would run on one Nvidia A100 GPU for fine-tuning. For the state space models, HyenaDNA, Mamba, and Caduceus, we pre-trained models using the pretraining script in the Caduceus github. Table 1 shows the model parameters that were chosen for each model. We have converted the perplexity values to a normalized unit of Bits per Character, which allows a more accurate comparison between models that use different tokenization methods.

We chose to use the Caduceus-ps (parameter-sharing) model to take full advantage of the new bi-mamba architecture that provides reverse compliment (RC) equivariance. The dimension parameter $d$ was fixed at 256 to correspond with the other 4-layer pretrained models in the Caduceus huggingface repository.

We pretrained the state space models with an input data length of 4096 nucleotides. This length was chosen because there are no publicly available pretrained HyenaDNA or Caduceus models with sequence lengths between 1k and 16k nucleotides. This input length made the input range comparable to that of the DNABERT2 and Nucleotide Transformer models, minimizing the influence of the inherent long-range capabilities of the state space models. This strategy allowed us to isolate tokenization, as much as possible when comparing very different pretrained models, as an experimental variable.

We used a grid search method of hyperparameter tuning, varying only the batch size and the learning rates for 10,000 steps. After determining the best hyperparameters for pre-training, we then trained the best model another 10,000 steps. The number of steps was chosen based on reported results in [10] and [14].

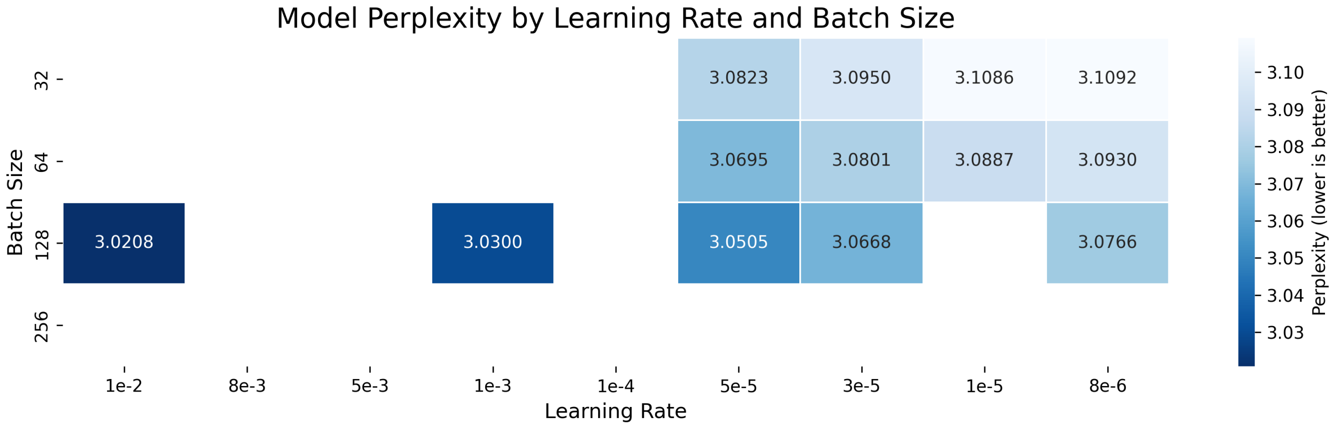


Fig. 1: Hyperparameter Tuning results for Pre-training the four layer Mamba model with byte-pair encoding. Missing cells are those that had NaN values. The model with the lowest perplexity after 10,000 steps was further trained until 20,000 steps and used as the model for fine-tuning. Similar plots were created for the other pretrained models

**Table 1.** Hyperparameter Tuning results for the models that we pretrained. Grid search hyperparameter tuning was completed for the first 10,000 steps of training and only the batch size and learning rate were varied. The model with the lowest perplexity was then further trained until 20,000 steps. The model parameters d, n, and k correspond to dimension, number of layers and input sequence length.

| | Fixed Parameters | | | | Hyperparameter Tuning | | Best HP | | Bits per Character |
|---|---|---|---|---|---|---|---|---|---|
| | d | n | k | RC Aug | Batch Size Range | Learning Rate Range | Batch Size | Learning Rate | Bits per Character |
| Mamba-char-4L | 256 | 4 | 4096 | TRUE | [128, 256] | [1e-2, 1e-3, 5e-3, 8e-3] | 256 | 1e-2 | 1.545 |
| Mamba-bpe-4L | 256 | 4 | 4096 | TRUE | [128, 256] | [1e-2, 8e-3, 5e-3, 1e-3, 1e-4, 5e-5, 3e-5, 1e-5, 8e-6] | 128 | 1e-2 | 0.304 |
| Caduceus-ps | 256 | 4 | 4096 | FALSE* | [128, 256] | [1e-3, 8e-3, 8e-4, 5e-4] | 256 | 1e-3 | 1.374 |

*RC Aug (Reverse Compliment Augmentation) was not necesary for the Caduceus-ps (parameter sharing) model because the reverse compliment sequences are passed through one half of the bi-Mamba architecture. Reverse Compliment augmentation was used in the Mamba models to make sure that the architecture saw the same data as Caduceus-ps.

To verify that we had achieved the best performance for each downstream task, we conducted a comprehensive grid-based hyperparameter search for every downstream fine-tuning task. The results of these hyperparameter tuning experiments can be seen in Tables 2, 3, 4, 5, and 6.

We observed significant instability when using higher learning rates and small batch sizes, with a high occurance of a NaN loss. In order to determine the frequency of the NaN outputs, we performed a larger analysis on the GUE dataset using 6 different learning rates and 4 different batch sizes. The results of this analysis can be seen in the appendix in Figure 2. Higher learning rates were associated with higher instability, but also had the highest accuracy. Lower learning rates were much more stable but needed to be trained much longer to obtain the same level of accuracy. Although our experiments were conducted on the Mamba-bpe model, this instability has also been observed by several other users using the character tokenized Caduceus model, as reported in the Caduceus Github repository. This instability seems to be related to initial seed values and can be resolved by reducing the learning-rate and/or changing seed values.

**Table 2.** Best hyperparameters for Genomic Benchmark benchmark

| Task | Model | LR | BS | MCC | Acc | N | MCC SD | Acc SD | LR Range | BS Range |
|---|---|---|---|---|---|---|---|---|---|---|
| demo_human_or_worm | Caduceus | 5.0e-04 | 128 | 0.943 | 0.972 | 10 | 0.0014 | 0.0007 | [1.0e-05, 1.0e-04, 5.0e-04, 1.0e-03, 2.0e-03] | [128, 256] |
| demo_human_or_worm | HyenaDNA | 6.0e-05 | 256 | 0.929 | 0.964 | 13 | 0.0031 | 0.0016 | [6.0e-05, 6.0e-04, 6.0e-03] | [128, 256] |
| demo_human_or_worm | Mamba-bpe | 1.0e-04 | 256 | 0.923 | 0.961 | 13 | 0.0024 | 0.0013 | [1.0e-05, 2.0e-05, 5.0e-05, 1.0e-04] | [128, 256] |
| demo_human_or_worm | Mamba-char | 1.0e-03 | 128 | 0.930 | 0.965 | 13 | 0.0086 | 0.0043 | [1.0e-04, 2.0e-04, 1.0e-03, 2.0e-03, 1.0e-02] | [128, 256] |
| dummy_mouse_enhancers | Caduceus | 2.0e-03 | 128 | 0.583 | 0.788 | 10 | 0.0377 | 0.0210 | [1.0e-05, 1.0e-04, 5.0e-04, 1.0e-03, 2.0e-03] | [128, 256] |
| dummy_mouse_enhancers | HyenaDNA | 6.0e-04 | 128 | 0.595 | 0.795 | 14 | 0.0487 | 0.0261 | [6.0e-05, 6.0e-04, 6.0e-03] | [128, 256] |
| dummy_mouse_enhancers | Mamba-bpe | 5.0e-05 | 256 | 0.555 | 0.775 | 13 | 0.0325 | 0.0186 | [1.0e-05, 5.0e-05, 1.0e-04] | [128, 256] |
| dummy_mouse_enhancers | Mamba-char | 1.0e-03 | 128 | 0.570 | 0.785 | 13 | 0.0393 | 0.0199 | [1.0e-03, 2.0e-03, 1.0e-02] | [128, 256] |
| human_enhancers_cohn | Caduceus | 1.0e-03 | 128 | 0.493 | 0.746 | 10 | 0.0078 | 0.0041 | [1.0e-05, 1.0e-04, 5.0e-04, 1.0e-03, 2.0e-03] | [128, 256] |
| human_enhancers_cohn | HyenaDNA | 6.0e-05 | 128 | 0.469 | 0.734 | 13 | 0.0077 | 0.0039 | [6.0e-05, 6.0e-04, 6.0e-03] | [128, 256] |
| human_enhancers_cohn | Mamba-bpe | 1.0e-05 | 256 | 0.478 | 0.739 | 13 | 0.0044 | 0.0023 | [1.0e-05, 2.0e-05, 5.0e-05, 1.0e-04] | [128, 256] |
| human_enhancers_cohn | Mamba-char | 1.0e-03 | 256 | 0.473 | 0.735 | 13 | 0.0100 | 0.0054 | [1.0e-04, 2.0e-04, 1.0e-03, 2.0e-03, 1.0e-02] | [128, 256] |
| human_enhancers_ensembl | Caduceus | 1.0e-03 | 128 | 0.822 | 0.911 | 10 | 0.0052 | 0.0027 | [1.0e-05, 1.0e-04, 5.0e-04, 1.0e-03, 2.0e-03] | [128, 256] |
| human_enhancers_ensembl | HyenaDNA | 6.0e-04 | 256 | 0.757 | 0.878 | 13 | 0.0055 | 0.0029 | [6.0e-05, 6.0e-04, 6.0e-03] | [128, 256] |
| human_enhancers_ensembl | Mamba-bpe | 1.0e-04 | 128 | 0.728 | 0.862 | 14 | 0.0314 | 0.0162 | [1.0e-05, 2.0e-05, 5.0e-05, 1.0e-04] | [128, 256] |
| human_enhancers_ensembl | Mamba-char | 1.0e-03 | 256 | 0.743 | 0.871 | 13 | 0.0505 | 0.0260 | [1.0e-04, 2.0e-04, 1.0e-03, 2.0e-03, 1.0e-02] | [128, 256] |
| human_ensembl_regulatory | Caduceus | 5.0e-04 | 128 | 0.807 | 0.869 | 6 | 0.0081 | 0.0070 | [1.0e-05, 1.0e-04, 5.0e-04, 1.0e-03, 2.0e-03] | [128, 256] |
| human_ensembl_regulatory | HyenaDNA | 6.0e-05 | 128 | 0.795 | 0.861 | 13 | 0.0061 | 0.0049 | [6.0e-05, 6.0e-04, 6.0e-03] | [128, 256] |
| human_ensembl_regulatory | Mamba-bpe | 1.0e-05 | 128 | 0.839 | 0.892 | 10 | 0.0026 | 0.0018 | [1.0e-05, 2.0e-05, 5.0e-05, 1.0e-04] | [128, 256] |
| human_ensembl_regulatory | Mamba-char | 1.0e-03 | 256 | 0.605 | 0.733 | 13 | 0.1359 | 0.0924 | [1.0e-04, 2.0e-04, 1.0e-03, 2.0e-03, 1.0e-02] | [128, 256] |
| human_nontata_promoters | Caduceus | 1.0e-03 | 256 | 0.891 | 0.945 | 10 | 0.0033 | 0.0015 | [1.0e-05, 1.0e-04, 5.0e-04, 1.0e-03, 2.0e-03] | [128, 256] |
| human_nontata_promoters | HyenaDNA | 6.0e-04 | 256 | 0.880 | 0.940 | 13 | 0.0072 | 0.0036 | [6.0e-05, 6.0e-04, 6.0e-03] | [128, 256] |
| human_nontata_promoters | Mamba-bpe | 1.0e-04 | 256 | 0.895 | 0.947 | 13 | 0.0038 | 0.0019 | [1.0e-05, 2.0e-05, 5.0e-05, 1.0e-04] | [128, 256] |
| human_nontata_promoters | Mamba-char | 1.0e-03 | 256 | 0.911 | 0.955 | 13 | 0.0082 | 0.0043 | [1.0e-04, 2.0e-04, 1.0e-03, 2.0e-03, 1.0e-02] | [128, 256] |
| human_ocr_ensembl | Caduceus | 1.0e-03 | 128 | 0.635 | 0.817 | 7 | 0.0062 | 0.0033 | [1.0e-05, 1.0e-04, 5.0e-04, 1.0e-03, 2.0e-03] | [128, 256] |
| human_ocr_ensembl | HyenaDNA | 6.0e-04 | 256 | 0.605 | 0.802 | 13 | 0.0037 | 0.0020 | [6.0e-05, 6.0e-04, 6.0e-03] | [128, 256] |
| human_ocr_ensembl | Mamba-bpe | 5.0e-05 | 128 | 0.545 | 0.772 | 13 | 0.0052 | 0.0026 | [1.0e-05, 2.0e-05, 5.0e-05, 1.0e-04] | [128, 256] |
| human_ocr_ensembl | Mamba-char | 2.0e-03 | 256 | 0.632 | 0.816 | 13 | 0.0035 | 0.0017 | [1.0e-04, 2.0e-04, 1.0e-03, 2.0e-03, 1.0e-02] | [128, 256] |

**Table 3.** Best hyperparameters for GUE benchmark (Part 1 of 2)

| Task | Model | LR | BS | MCC | Acc | N | MCC SD | Acc SD | LR Range | BS Range |
|---|---|---|---|---|---|---|---|---|---|---|
| covid | Caduceus | 3.0e-04 | 128 | 0.630 | 0.672 | 4 | 0.0150 | 0.0135 | [8.0e-05, 1.0e-04, 3.0e-04, 5.0e-04, 8.0e-04, 1.0e-03, 2.0e-03] | [128, 256, 512] |
| covid | HyenaDNA | 6.0e-05 | 128 | 0.620 | 0.663 | 11 | 0.0205 | 0.0180 | [6.0e-05, 6.0e-04, 6.0e-03] | [128, 256] |
| covid | Mamba-bpe | 2.0e-04 | 128 | 0.652 | 0.691 | 13 | 0.0243 | 0.0232 | [5.0e-05, 1.0e-04, 2.0e-04, 1.0e-03] | [128, 256] |
| covid | Mamba-char | 2.0e-04 | 128 | 0.576 | 0.625 | 10 | 0.0110 | 0.0097 | [5.0e-05, 1.0e-04, 2.0e-04] | [128, 256] |
| mouse_0 | Caduceus | 2.0e-03 | 128 | 0.610 | 0.803 | 10 | 0.0151 | 0.0080 | [1.0e-05, 1.0e-04, 5.0e-04, 1.0e-03, 2.0e-03] | [128, 256] |
| mouse_0 | HyenaDNA | 6.0e-04 | 256 | 0.523 | 0.760 | 10 | 0.0220 | 0.0129 | [6.0e-05, 6.0e-04, 6.0e-03] | [128, 256] |
| mouse_0 | Mamba-bpe | 1.0e-04 | 128 | 0.507 | 0.752 | 13 | 0.0161 | 0.0082 | [1.0e-05, 2.0e-05, 5.0e-05, 1.0e-04, 2.0e-04, 6.0e-04] | [128, 256] |
| mouse_0 | Mamba-char | 2.0e-04 | 128 | 0.465 | 0.731 | 10 | 0.0185 | 0.0097 | [5.0e-05, 1.0e-04, 2.0e-04] | [128, 256] |
| mouse_1 | Caduceus | 1.0e-03 | 128 | 0.845 | 0.923 | 10 | 0.0048 | 0.0025 | [1.0e-05, 1.0e-04, 5.0e-04, 1.0e-03, 2.0e-03] | [128, 256] |
| mouse_1 | HyenaDNA | 6.0e-04 | 256 | 0.824 | 0.911 | 10 | 0.0036 | 0.0020 | [6.0e-05, 6.0e-04, 6.0e-03] | [128, 256] |
| mouse_1 | Mamba-bpe | 2.0e-04 | 256 | 0.801 | 0.900 | 13 | 0.0055 | 0.0027 | [1.0e-05, 2.0e-05, 5.0e-05, 1.0e-04, 2.0e-04] | [128, 256] |
| mouse_1 | Mamba-char | 2.0e-04 | 256 | 0.795 | 0.897 | 10 | 0.0035 | 0.0016 | [5.0e-05, 1.0e-04, 2.0e-04] | [128, 256] |
| mouse_2 | Caduceus | 2.0e-03 | 128 | 0.845 | 0.921 | 7 | 0.0167 | 0.0081 | [1.0e-05, 1.0e-04, 2.0e-04, 5.0e-04, 1.0e-03, 2.0e-03] | [128, 256] |
| mouse_2 | HyenaDNA | 6.0e-04 | 128 | 0.801 | 0.899 | 10 | 0.0320 | 0.0162 | [6.0e-05, 6.0e-04, 6.0e-03] | [128, 256] |
| mouse_2 | Mamba-bpe | 2.0e-04 | 128 | 0.772 | 0.885 | 13 | 0.0212 | 0.0107 | [1.0e-05, 2.0e-05, 5.0e-05, 1.0e-04, 2.0e-04, 2.0e-03] | [128, 256] |
| mouse_2 | Mamba-char | 2.0e-04 | 128 | 0.701 | 0.849 | 17 | 0.0274 | 0.0135 | [1.0e-04, 2.0e-04] | [128, 256] |
| mouse_3 | Caduceus | 2.0e-03 | 128 | 0.827 | 0.911 | 10 | 0.0215 | 0.0108 | [1.0e-05, 1.0e-04, 5.0e-04, 1.0e-03, 2.0e-03] | [128, 256] |
| mouse_3 | HyenaDNA | 6.0e-04 | 128 | 0.737 | 0.866 | 10 | 0.0477 | 0.0227 | [6.0e-05, 6.0e-04, 6.0e-03] | [128, 256] |
| mouse_3 | Mamba-bpe | 2.0e-04 | 128 | 0.588 | 0.792 | 13 | 0.0471 | 0.0246 | [1.0e-05, 2.0e-05, 5.0e-05, 1.0e-04, 2.0e-04, 2.0e-03] | [128, 256] |
| mouse_3 | Mamba-char | 2.0e-04 | 128 | 0.501 | 0.741 | 10 | 0.0283 | 0.0131 | [5.0e-05, 1.0e-04, 2.0e-04] | [128, 256] |
| mouse_4 | Caduceus | 2.0e-03 | 256 | 0.496 | 0.746 | 10 | 0.0075 | 0.0045 | [1.0e-05, 1.0e-04, 5.0e-04, 1.0e-03, 2.0e-03] | [128, 256] |
| mouse_4 | HyenaDNA | 6.0e-04 | 128 | 0.468 | 0.733 | 10 | 0.0159 | 0.0080 | [6.0e-05, 6.0e-04, 6.0e-03] | [128, 256] |
| mouse_4 | Mamba-bpe | 5.0e-05 | 128 | 0.418 | 0.709 | 13 | 0.0128 | 0.0067 | [1.0e-05, 2.0e-05, 5.0e-05, 1.0e-04, 2.0e-04] | [128, 256] |
| mouse_4 | Mamba-char | 2.0e-04 | 128 | 0.423 | 0.711 | 10 | 0.0165 | 0.0079 | [5.0e-05, 1.0e-04, 2.0e-04] | [128, 256] |
| prom_300_all | Caduceus | 5.0e-04 | 128 | 0.821 | 0.910 | 10 | 0.0054 | 0.0026 | [1.0e-05, 1.0e-04, 5.0e-04, 1.0e-03, 2.0e-03] | [128, 256] |
| prom_300_all | HyenaDNA | 6.0e-04 | 128 | 0.826 | 0.912 | 10 | 0.0052 | 0.0026 | [6.0e-05, 6.0e-04, 6.0e-03] | [128, 256] |
| prom_300_all | Mamba-bpe | 5.0e-05 | 256 | 0.810 | 0.904 | 13 | 0.0032 | 0.0016 | [1.0e-05, 2.0e-05, 5.0e-05, 1.0e-04, 2.0e-04] | [128, 256] |
| prom_300_all | Mamba-char | 1.0e-04 | 128 | 0.818 | 0.909 | 10 | 0.0065 | 0.0031 | [5.0e-05, 1.0e-04, 2.0e-04] | [128, 256] |
| prom_300_notata | Caduceus | 1.0e-03 | 128 | 0.928 | 0.964 | 10 | 0.0021 | 0.0011 | [1.0e-05, 1.0e-04, 5.0e-04, 1.0e-03, 2.0e-03] | [128, 256] |
| prom_300_notata | HyenaDNA | 6.0e-04 | 256 | 0.918 | 0.959 | 10 | 0.0054 | 0.0027 | [6.0e-05, 6.0e-04, 6.0e-03] | [128, 256] |
| prom_300_notata | Mamba-bpe | 5.0e-05 | 128 | 0.911 | 0.955 | 13 | 0.0046 | 0.0024 | [1.0e-05, 2.0e-05, 5.0e-05, 1.0e-04, 2.0e-04] | [128, 256] |
| prom_300_notata | Mamba-char | 2.0e-04 | 128 | 0.917 | 0.958 | 10 | 0.0027 | 0.0014 | [5.0e-05, 1.0e-04, 2.0e-04] | [128, 256] |
| prom_300_tata | Caduceus | 2.0e-03 | 128 | 0.581 | 0.789 | 10 | 0.0317 | 0.0162 | [1.0e-05, 1.0e-04, 5.0e-04, 1.0e-03, 2.0e-03] | [128, 256] |
| prom_300_tata | HyenaDNA | 6.0e-04 | 128 | 0.466 | 0.732 | 10 | 0.0431 | 0.0221 | [6.0e-05, 6.0e-04, 6.0e-03] | [128, 256] |
| prom_300_tata | Mamba-bpe | 5.0e-05 | 128 | 0.310 | 0.655 | 13 | 0.0330 | 0.0156 | [1.0e-05, 2.0e-05, 5.0e-05, 1.0e-04, 2.0e-04] | [128, 256] |
| prom_300_tata | Mamba-char | 2.0e-04 | 128 | 0.334 | 0.667 | 10 | 0.0300 | 0.0158 | [5.0e-05, 1.0e-04, 2.0e-04] | [128, 256] |
| prom_core_all | Caduceus | 1.0e-03 | 256 | 0.656 | 0.828 | 10 | 0.0085 | 0.0040 | [1.0e-05, 1.0e-04, 5.0e-04, 1.0e-03, 2.0e-03] | [128, 256] |

**Table 4.** Best hyperparameters for GUE benchmark (Part 2 of 2)

| Task | Model | LR | BS | MCC | Acc | N | MCC SD | Acc SD | LR Range | BS Range |
|---|---|---|---|---|---|---|---|---|---|---|
| prom_core_all | HyenaDNA | 6.0e-04 | 256 | 0.626 | 0.813 | 10 | 0.0077 | 0.0039 | [6.0e-05, 6.0e-04, 6.0e-03] | [128, 256] |
| prom_core_all | Mamba-bpe | 1.0e-04 | 128 | 0.596 | 0.798 | 13 | 0.0089 | 0.0044 | [1.0e-05, 2.0e-05, 5.0e-05, 1.0e-04, 2.0e-04, 5.0e-04] | [128, 256] |
| prom_core_all | Mamba-char | 2.0e-04 | 128 | 0.617 | 0.808 | 10 | 0.0069 | 0.0034 | [5.0e-05, 1.0e-04, 2.0e-04] | [128, 256] |
| prom_core_notata | Caduceus | 2.0e-03 | 256 | 0.675 | 0.836 | 10 | 0.0074 | 0.0047 | [1.0e-05, 1.0e-04, 5.0e-04, 1.0e-03, 2.0e-03] | [128, 256] |
| prom_core_notata | HyenaDNA | 6.0e-04 | 256 | 0.664 | 0.832 | 10 | 0.0036 | 0.0018 | [6.0e-05, 6.0e-04, 6.0e-03] | [128, 256] |
| prom_core_notata | Mamba-bpe | 5.0e-05 | 256 | 0.652 | 0.826 | 13 | 0.0066 | 0.0033 | [1.0e-05, 2.0e-05, 5.0e-05, 1.0e-04, 2.0e-04] | [128, 256] |
| prom_core_notata | Mamba-char | 2.0e-04 | 128 | 0.654 | 0.827 | 10 | 0.0042 | 0.0021 | [5.0e-05, 1.0e-04, 2.0e-04] | [128, 256] |
| prom_core_tata | Caduceus | 2.0e-03 | 128 | 0.595 | 0.794 | 10 | 0.1272 | 0.0675 | [1.0e-05, 1.0e-04, 2.0e-04, 5.0e-04, 1.0e-03, 2.0e-03] | [128, 256] |
| prom_core_tata | HyenaDNA | 6.0e-04 | 128 | 0.465 | 0.732 | 11 | 0.0314 | 0.0157 | [6.0e-05, 6.0e-04, 6.0e-03] | [128, 256] |
| prom_core_tata | Mamba-bpe | 2.0e-04 | 256 | 0.424 | 0.711 | 13 | 0.0279 | 0.0142 | [1.0e-05, 2.0e-05, 5.0e-05, 1.0e-04, 2.0e-04] | [128, 256] |
| prom_core_tata | Mamba-char | 2.0e-04 | 128 | 0.540 | 0.769 | 18 | 0.0177 | 0.0090 | [5.0e-05, 1.0e-04, 2.0e-04] | [128, 256] |
| reconstructed | Caduceus | 1.0e-03 | 256 | 0.816 | 0.893 | 10 | 0.0248 | 0.0144 | [1.0e-04, 1.0e-03, 2.0e-03] | [128, 256] |
| reconstructed | HyenaDNA | 6.0e-04 | 128 | 0.820 | 0.895 | 10 | 0.0119 | 0.0067 | [6.0e-05, 6.0e-04, 6.0e-03] | [128, 256] |
| reconstructed | Mamba-bpe | 2.0e-04 | 128 | 0.614 | 0.776 | 13 | 0.0157 | 0.0088 | [5.0e-05, 1.0e-04, 2.0e-04] | [128, 256] |
| reconstructed | Mamba-char | 2.0e-04 | 128 | 0.664 | 0.804 | 10 | 0.0249 | 0.0151 | [5.0e-05, 1.0e-04, 2.0e-04] | [128, 256] |
| tf_0 | Caduceus | 5.0e-04 | 256 | 0.664 | 0.826 | 10 | 0.0134 | 0.0087 | [1.0e-05, 1.0e-04, 5.0e-04, 1.0e-03, 2.0e-03] | [128, 256] |
| tf_0 | HyenaDNA | 6.0e-05 | 256 | 0.671 | 0.830 | 12 | 0.0082 | 0.0055 | [6.0e-05, 6.0e-04, 6.0e-03] | [128, 256] |
| tf_0 | Mamba-bpe | 1.0e-04 | 256 | 0.643 | 0.815 | 13 | 0.0165 | 0.0100 | [5.0e-05, 1.0e-04, 2.0e-04, 1.0e-03] | [128, 256] |
| tf_0 | Mamba-char | 2.0e-04 | 256 | 0.671 | 0.830 | 10 | 0.0128 | 0.0067 | [5.0e-05, 1.0e-04, 2.0e-04] | [128, 256] |
| tf_1 | Caduceus | 1.0e-03 | 128 | 0.716 | 0.852 | 10 | 0.0180 | 0.0095 | [1.0e-05, 1.0e-04, 5.0e-04, 1.0e-03, 2.0e-03] | [128, 256] |
| tf_1 | HyenaDNA | 6.0e-05 | 256 | 0.706 | 0.848 | 12 | 0.0123 | 0.0088 | [6.0e-05, 6.0e-04, 6.0e-03] | [128, 256] |
| tf_1 | Mamba-bpe | 2.0e-04 | 128 | 0.683 | 0.836 | 12 | 0.0154 | 0.0107 | [5.0e-05, 1.0e-04, 2.0e-04, 6.0e-04, 2.0e-03] | [128, 256] |
| tf_1 | Mamba-char | 5.0e-05 | 128 | 0.691 | 0.840 | 10 | 0.0094 | 0.0052 | [5.0e-05, 1.0e-04, 2.0e-04] | [128, 256] |
| tf_2 | Caduceus | 1.0e-03 | 128 | 0.705 | 0.847 | 10 | 0.0331 | 0.0188 | [1.0e-05, 1.0e-04, 5.0e-04, 1.0e-03, 2.0e-03] | [128, 256] |
| tf_2 | HyenaDNA | 6.0e-04 | 128 | 0.671 | 0.833 | 11 | 0.0224 | 0.0113 | [6.0e-05, 6.0e-04, 6.0e-03] | [128, 256] |
| tf_2 | Mamba-bpe | 1.0e-04 | 128 | 0.571 | 0.782 | 13 | 0.0140 | 0.0067 | [5.0e-05, 1.0e-04, 2.0e-04] | [128, 256] |
| tf_2 | Mamba-char | 2.0e-04 | 128 | 0.613 | 0.802 | 10 | 0.0243 | 0.0137 | [5.0e-05, 1.0e-04, 2.0e-04] | [128, 256] |
| tf_3 | Caduceus | 2.0e-03 | 128 | 0.539 | 0.766 | 10 | 0.0544 | 0.0275 | [1.0e-05, 1.0e-04, 5.0e-04, 1.0e-03, 2.0e-03] | [128, 256] |
| tf_3 | HyenaDNA | 6.0e-04 | 128 | 0.519 | 0.757 | 11 | 0.0195 | 0.0100 | [6.0e-05, 6.0e-04, 6.0e-03] | [128, 256] |
| tf_3 | Mamba-bpe | 1.0e-04 | 128 | 0.480 | 0.736 | 13 | 0.0169 | 0.0110 | [5.0e-05, 1.0e-04, 2.0e-04, 6.0e-04] | [128, 256] |
| tf_3 | Mamba-char | 1.0e-04 | 128 | 0.485 | 0.739 | 10 | 0.0217 | 0.0107 | [5.0e-05, 1.0e-04, 2.0e-04] | [128, 256] |
| tf_4 | Caduceus | 2.0e-03 | 128 | 0.786 | 0.892 | 10 | 0.0307 | 0.0153 | [1.0e-05, 1.0e-04, 5.0e-04, 1.0e-03, 2.0e-03] | [128, 256] |
| tf_4 | HyenaDNA | 6.0e-04 | 128 | 0.749 | 0.874 | 10 | 0.0158 | 0.0079 | [6.0e-05, 6.0e-04, 6.0e-03] | [128, 256] |
| tf_4 | Mamba-bpe | 2.0e-04 | 128 | 0.697 | 0.848 | 13 | 0.0208 | 0.0104 | [5.0e-05, 1.0e-04, 2.0e-04, 6.0e-04] | [128, 256] |
| tf_4 | Mamba-char | 1.0e-04 | 256 | 0.706 | 0.851 | 10 | 0.0060 | 0.0031 | [5.0e-05, 1.0e-04, 2.0e-04] | [128, 256] |

**Table 5.** Best hyperparameters for Nucleotide Transformer (revised) benchmark (Part 1 of 2)

| Task | Model | LR | BS | MCC | Acc | N | MCC SD | Acc SD | LR Range | BS Range |
|---|---|---|---|---|---|---|---|---|---|---|
| H2AFZ | Caduceus | 1.0e-04 | 128 | 0.432 | 0.708 | 7 | 0.0035 | 0.0012 | [1.0e-05, 1.0e-04, 1.0e-03, 2.0e-03, 8.0e-03, 1.0e-02] | [128, 256] |
| H2AFZ | HyenaDNA | 6.0e-04 | 128 | 0.413 | 0.702 | 13 | 0.0073 | 0.0035 | [6.0e-05, 2.0e-04, 6.0e-04, 6.0e-03] | [128, 256] |
| H2AFZ | Mamba-bpe | 5.0e-05 | 128 | 0.405 | 0.693 | 13 | 0.0128 | 0.0089 | [5.0e-05, 1.0e-04, 2.0e-04] | [128, 256] |
| H2AFZ | Mamba-char | 5.0e-04 | 128 | 0.392 | 0.688 | 18 | 0.0149 | 0.0107 | [1.0e-04, 3.0e-04, 5.0e-04, 1.0e-03, 2.0e-03, 1.0e-02] | [128, 256] |
| H3K27ac | Caduceus | 1.0e-03 | 128 | 0.467 | 0.733 | 3 | 0.0323 | 0.0162 | [1.0e-05, 1.0e-04, 1.0e-03, 2.0e-03, 8.0e-03, 1.0e-02] | [128, 256] |
| H3K27ac | HyenaDNA | 6.0e-05 | 256 | 0.421 | 0.707 | 13 | 0.0133 | 0.0078 | [6.0e-05, 2.0e-04, 6.0e-04, 6.0e-03] | [128, 256] |
| H3K27ac | Mamba-bpe | 5.0e-05 | 256 | 0.392 | 0.693 | 14 | 0.0145 | 0.0055 | [5.0e-05, 1.0e-04, 2.0e-04] | [128, 256] |
| H3K27ac | Mamba-char | 1.0e-04 | 256 | 0.418 | 0.705 | 18 | 0.0125 | 0.0101 | [1.0e-04, 3.0e-04, 5.0e-04, 1.0e-03, 2.0e-03, 1.0e-02] | [128, 256] |
| H3K27me3 | Caduceus | 1.0e-03 | 128 | 0.551 | 0.763 | 3 | 0.0072 | 0.0047 | [1.0e-05, 1.0e-04, 1.0e-03, 2.0e-03, 8.0e-03, 1.0e-02] | [128, 256] |
| H3K27me3 | HyenaDNA | 6.0e-05 | 256 | 0.542 | 0.764 | 13 | 0.0133 | 0.0078 | [6.0e-05, 2.0e-04, 6.0e-04, 6.0e-03] | [128, 256] |
| H3K27me3 | Mamba-bpe | 5.0e-05 | 128 | 0.530 | 0.752 | 13 | 0.0096 | 0.0068 | [5.0e-05, 1.0e-04, 2.0e-04] | [128, 256] |
| H3K27me3 | Mamba-char | 5.0e-04 | 128 | 0.535 | 0.757 | 18 | 0.0105 | 0.0083 | [1.0e-04, 3.0e-04, 5.0e-04, 1.0e-03, 2.0e-03, 1.0e-02] | [128, 256] |
| H3K36me3 | Caduceus | 1.0e-03 | 256 | 0.531 | 0.760 | 3 | 0.0179 | 0.0076 | [1.0e-05, 1.0e-04, 1.0e-03, 2.0e-03, 8.0e-03, 1.0e-02] | [128, 256] |
| H3K36me3 | HyenaDNA | 6.0e-04 | 256 | 0.504 | 0.746 | 13 | 0.0079 | 0.0059 | [6.0e-05, 2.0e-04, 6.0e-04, 6.0e-03] | [128, 256] |
| H3K36me3 | Mamba-bpe | 5.0e-05 | 128 | 0.489 | 0.738 | 13 | 0.0096 | 0.0058 | [5.0e-05, 1.0e-04, 2.0e-04, 3.0e-04] | [128, 256] |
| H3K36me3 | Mamba-char | 3.0e-04 | 128 | 0.486 | 0.737 | 18 | 0.0122 | 0.0088 | [1.0e-04, 3.0e-04, 5.0e-04, 1.0e-03, 2.0e-03, 1.0e-02] | [128, 256] |
| H3K4me1 | Caduceus | 1.0e-03 | 128 | 0.429 | 0.706 | 11 | 0.0082 | 0.0101 | [1.0e-05, 1.0e-04, 1.0e-03, 2.0e-03, 8.0e-03, 1.0e-02] | [128, 256] |
| H3K4me1 | HyenaDNA | 6.0e-04 | 128 | 0.403 | 0.695 | 11 | 0.0074 | 0.0041 | [6.0e-05, 2.0e-04, 6.0e-04, 6.0e-03] | [128, 256] |
| H3K4me1 | Mamba-bpe | 5.0e-05 | 128 | 0.406 | 0.693 | 13 | 0.0095 | 0.0053 | [5.0e-05, 1.0e-04, 2.0e-04] | [128, 256] |
| H3K4me1 | Mamba-char | 1.0e-04 | 128 | 0.405 | 0.692 | 18 | 0.0101 | 0.0054 | [1.0e-04, 3.0e-04, 5.0e-04, 1.0e-03, 2.0e-03, 1.0e-02] | [128, 256] |
| H3K4me2 | Caduceus | 1.0e-03 | 128 | 0.485 | 0.736 | 7 | 0.0196 | 0.0121 | [1.0e-05, 1.0e-04, 1.0e-03, 2.0e-03, 8.0e-03, 1.0e-02] | [128, 256] |
| H3K4me2 | HyenaDNA | 6.0e-04 | 128 | 0.477 | 0.732 | 13 | 0.0146 | 0.0091 | [6.0e-05, 2.0e-04, 6.0e-04, 6.0e-03] | [128, 256] |
| H3K4me2 | Mamba-bpe | 5.0e-05 | 256 | 0.470 | 0.729 | 14 | 0.0144 | 0.0103 | [5.0e-05, 1.0e-04, 2.0e-04] | [128, 256] |
| H3K4me2 | Mamba-char | 1.0e-04 | 256 | 0.456 | 0.724 | 18 | 0.0143 | 0.0080 | [1.0e-04, 3.0e-04, 5.0e-04, 1.0e-03, 2.0e-03, 1.0e-02] | [128, 256] |
| H3K4me3 | Caduceus | 1.0e-04 | 128 | 0.623 | 0.811 | 10 | 0.0086 | 0.0045 | [1.0e-05, 1.0e-04, 1.0e-03, 2.0e-03, 8.0e-03, 1.0e-02] | [128, 256] |
| H3K4me3 | HyenaDNA | 6.0e-04 | 256 | 0.600 | 0.799 | 11 | 0.0127 | 0.0057 | [6.0e-05, 2.0e-04, 6.0e-04, 6.0e-03] | [128, 256] |
| H3K4me3 | Mamba-bpe | 5.0e-05 | 256 | 0.601 | 0.800 | 13 | 0.0158 | 0.0080 | [5.0e-05, 1.0e-04, 2.0e-04] | [128, 256] |
| H3K4me3 | Mamba-char | 1.0e-04 | 256 | 0.606 | 0.802 | 18 | 0.0137 | 0.0073 | [1.0e-04, 3.0e-04, 5.0e-04, 1.0e-03, 2.0e-03, 1.0e-02] | [128, 256] |
| H3K9ac | Caduceus | 1.0e-04 | 256 | 0.512 | 0.755 | 3 | 0.0039 | 0.0023 | [1.0e-05, 1.0e-04, 1.0e-03, 2.0e-03, 8.0e-03, 1.0e-02] | [128, 256] |
| H3K9ac | HyenaDNA | 2.0e-04 | 256 | 0.505 | 0.751 | 12 | 0.0117 | 0.0068 | [6.0e-05, 2.0e-04, 6.0e-04, 6.0e-03] | [128, 256] |
| H3K9ac | Mamba-bpe | 5.0e-05 | 128 | 0.477 | 0.737 | 13 | 0.0212 | 0.0103 | [5.0e-05, 1.0e-04, 2.0e-04] | [128, 256] |
| H3K9ac | Mamba-char | 5.0e-04 | 128 | 0.502 | 0.749 | 18 | 0.0193 | 0.0101 | [1.0e-04, 3.0e-04, 5.0e-04, 1.0e-03, 2.0e-03, 1.0e-02] | [128, 256] |
| H3K9me3 | Caduceus | 1.0e-03 | 256 | 0.395 | 0.697 | 3 | 0.0215 | 0.0105 | [1.0e-05, 1.0e-04, 1.0e-03, 2.0e-03, 8.0e-03, 1.0e-02] | [128, 256] |
| H3K9me3 | HyenaDNA | 6.0e-04 | 256 | 0.370 | 0.681 | 13 | 0.0308 | 0.0164 | [6.0e-05, 2.0e-04, 6.0e-04, 6.0e-03] | [128, 256] |
| H3K9me3 | Mamba-bpe | 1.0e-04 | 128 | 0.290 | 0.643 | 13 | 0.0285 | 0.0144 | [5.0e-05, 1.0e-04, 2.0e-04] | [128, 256] |
| H3K9me3 | Mamba-char | 3.0e-04 | 128 | 0.340 | 0.668 | 18 | 0.0168 | 0.0089 | [1.0e-04, 3.0e-04, 5.0e-04, 1.0e-03, 2.0e-03, 1.0e-02] | [128, 256] |
| H4K20me1 | Caduceus | 1.0e-03 | 128 | 0.589 | 0.788 | 3 | 0.0059 | 0.0024 | [1.0e-05, 1.0e-04, 1.0e-03, 2.0e-03, 8.0e-03, 1.0e-02] | [128, 256] |

**Table 6.** Best hyperparameters for Nucleotide Transformer (revised) benchmark (Part 2 of 2)

| Task | Model | LR | BS | MCC | Acc | N | MCC SD | Acc SD | LR Range | BS Range |
|---|---|---|---|---|---|---|---|---|---|---|
| H4K20me1 | HyenaDNA | 6.0e-05 | 256 | 0.574 | 0.782 | 13 | 0.0137 | 0.0086 | [6.0e-05, 2.0e-04, 6.0e-04, 6.0e-03] | [128, 256] |
| H4K20me1 | Mamba-bpe | 5.0e-05 | 256 | 0.544 | 0.766 | 14 | 0.0090 | 0.0066 | [5.0e-05, 1.0e-04, 2.0e-04] | [128, 256] |
| H4K20me1 | Mamba-char | 5.0e-04 | 256 | 0.560 | 0.775 | 18 | 0.0105 | 0.0085 | [1.0e-04, 3.0e-04, 5.0e-04, 1.0e-03, 2.0e-03, 1.0e-02] | [128, 256] |
| enhancers | Caduceus | 1.0e-03 | 128 | 0.505 | 0.745 | 10 | 0.0083 | 0.0082 | [1.0e-05, 1.0e-04, 1.0e-03, 2.0e-03, 8.0e-03, 1.0e-02] | [128, 256] |
| enhancers | HyenaDNA | 2.0e-04 | 256 | 0.492 | 0.740 | 13 | 0.0108 | 0.0044 | [6.0e-05, 2.0e-04, 6.0e-04, 6.0e-03] | [128, 256] |
| enhancers | Mamba-bpe | 5.0e-05 | 256 | 0.487 | 0.736 | 13 | 0.0079 | 0.0057 | [5.0e-05, 1.0e-04, 2.0e-04] | [128, 256] |
| enhancers | Mamba-char | 2.0e-03 | 256 | 0.489 | 0.737 | 13 | 0.0111 | 0.0049 | [1.0e-03, 2.0e-03, 1.0e-02] | [128, 256] |
| enhancers_types | Caduceus | 1.0e-03 | 128 | 0.461 | 0.709 | 10 | 0.0101 | 0.0110 | [1.0e-05, 1.0e-04, 1.0e-03, 2.0e-03, 8.0e-03, 1.0e-02] | [128, 256] |
| enhancers_types | HyenaDNA | 2.0e-04 | 256 | 0.447 | 0.703 | 11 | 0.0114 | 0.0065 | [6.0e-05, 2.0e-04, 6.0e-04, 6.0e-03] | [128, 256] |
| enhancers_types | Mamba-bpe | 5.0e-05 | 256 | 0.449 | 0.700 | 13 | 0.0101 | 0.0056 | [5.0e-05, 1.0e-04, 2.0e-04] | [128, 256] |
| enhancers_types | Mamba-char | 2.0e-03 | 128 | 0.462 | 0.709 | 18 | 0.0091 | 0.0075 | [1.0e-03, 2.0e-03, 1.0e-02] | [128, 256] |
| promoter_all | Caduceus | 1.0e-05 | 128 | 0.718 | 0.859 | 10 | 0.0045 | 0.0021 | [1.0e-05, 1.0e-04, 1.0e-03, 2.0e-03, 8.0e-03, 1.0e-02] | [128, 256] |
| promoter_all | HyenaDNA | 6.0e-05 | 128 | 0.708 | 0.854 | 11 | 0.0102 | 0.0051 | [6.0e-05, 2.0e-04, 6.0e-04, 6.0e-03] | [128, 256] |
| promoter_all | Mamba-bpe | 2.0e-04 | 128 | 0.698 | 0.848 | 13 | 0.0112 | 0.0056 | [5.0e-05, 1.0e-04, 2.0e-04] | [128, 256] |
| promoter_all | Mamba-char | 2.0e-03 | 128 | 0.711 | 0.854 | 13 | 0.0086 | 0.0043 | [1.0e-03, 2.0e-03, 1.0e-02] | [128, 256] |
| promoter_no_tata | Caduceus | 1.0e-04 | 128 | 0.738 | 0.868 | 10 | 0.0081 | 0.0046 | [1.0e-05, 1.0e-04, 1.0e-03, 2.0e-03, 8.0e-03, 1.0e-02] | [128, 256] |
| promoter_no_tata | HyenaDNA | 6.0e-05 | 256 | 0.735 | 0.867 | 13 | 0.0052 | 0.0030 | [6.0e-05, 2.0e-04, 6.0e-04, 6.0e-03] | [128, 256] |
| promoter_no_tata | Mamba-bpe | 5.0e-05 | 256 | 0.727 | 0.863 | 13 | 0.0086 | 0.0043 | [5.0e-05, 1.0e-04, 2.0e-04] | [128, 256] |
| promoter_no_tata | Mamba-char | 2.0e-03 | 128 | 0.733 | 0.866 | 13 | 0.0097 | 0.0048 | [1.0e-03, 2.0e-03, 1.0e-02] | [128, 256] |
| promoter_tata | Caduceus | 1.0e-03 | 128 | 0.838 | 0.919 | 10 | 0.0347 | 0.0173 | [1.0e-05, 1.0e-04, 1.0e-03, 2.0e-03, 8.0e-03, 1.0e-02] | [128, 256] |
| promoter_tata | HyenaDNA | 2.0e-04 | 128 | 0.671 | 0.834 | 11 | 0.0328 | 0.0164 | [6.0e-05, 2.0e-04, 6.0e-04, 6.0e-03] | [128, 256] |
| promoter_tata | Mamba-bpe | 2.0e-04 | 256 | 0.621 | 0.808 | 13 | 0.0318 | 0.0168 | [5.0e-05, 1.0e-04, 2.0e-04] | [128, 256] |
| promoter_tata | Mamba-char | 1.0e-03 | 128 | 0.676 | 0.836 | 13 | 0.0403 | 0.0207 | [1.0e-03, 2.0e-03, 1.0e-02] | [128, 256] |
| splice_sites_acceptors | Caduceus | 1.0e-03 | 128 | 0.873 | 0.937 | 10 | 0.0090 | 0.0045 | [1.0e-05, 1.0e-04, 1.0e-03, 2.0e-03, 8.0e-03, 1.0e-02] | [128, 256] |
| splice_sites_acceptors | HyenaDNA | 6.0e-04 | 128 | 0.887 | 0.944 | 13 | 0.0164 | 0.0081 | [6.0e-05, 2.0e-04, 6.0e-04, 6.0e-03] | [128, 256] |
| splice_sites_acceptors | Mamba-bpe | 2.0e-04 | 128 | 0.607 | 0.802 | 13 | 0.0290 | 0.0159 | [5.0e-05, 1.0e-04, 2.0e-04] | [128, 256] |
| splice_sites_acceptors | Mamba-char | 2.0e-03 | 128 | 0.902 | 0.951 | 13 | 0.0072 | 0.0036 | [1.0e-03, 2.0e-03, 1.0e-02] | [128, 256] |
| splice_sites_all | Caduceus | 1.0e-03 | 128 | 0.911 | 0.941 | 10 | 0.0102 | 0.0068 | [1.0e-05, 1.0e-04, 1.0e-03, 2.0e-03, 8.0e-03, 1.0e-02] | [128, 256] |
| splice_sites_all | HyenaDNA | 6.0e-04 | 128 | 0.909 | 0.939 | 13 | 0.0155 | 0.0103 | [6.0e-05, 2.0e-04, 6.0e-04, 6.0e-03] | [128, 256] |
| splice_sites_all | Mamba-bpe | 2.0e-04 | 128 | 0.705 | 0.803 | 13 | 0.0126 | 0.0089 | [5.0e-05, 1.0e-04, 2.0e-04] | [128, 256] |
| splice_sites_all | Mamba-char | 2.0e-03 | 128 | 0.930 | 0.953 | 13 | 0.0057 | 0.0039 | [1.0e-03, 2.0e-03, 1.0e-02] | [128, 256] |
| splice_sites_donors | Caduceus | 1.0e-03 | 256 | 0.880 | 0.940 | 3 | 0.0164 | 0.0082 | [1.0e-05, 1.0e-04, 1.0e-03, 2.0e-03, 8.0e-03, 1.0e-02] | [128, 256] |
| splice_sites_donors | HyenaDNA | 6.0e-04 | 128 | 0.919 | 0.960 | 13 | 0.0305 | 0.0153 | [6.0e-05, 2.0e-04, 6.0e-04, 6.0e-03] | [128, 256] |
| splice_sites_donors | Mamba-bpe | 2.0e-04 | 128 | 0.598 | 0.797 | 13 | 0.0173 | 0.0096 | [5.0e-05, 1.0e-04, 2.0e-04] | [128, 256] |
| splice_sites_donors | Mamba-char | 2.0e-03 | 128 | 0.919 | 0.960 | 18 | 0.0534 | 0.0268 | [1.0e-03, 2.0e-03, 1.0e-02] | [128, 256] |

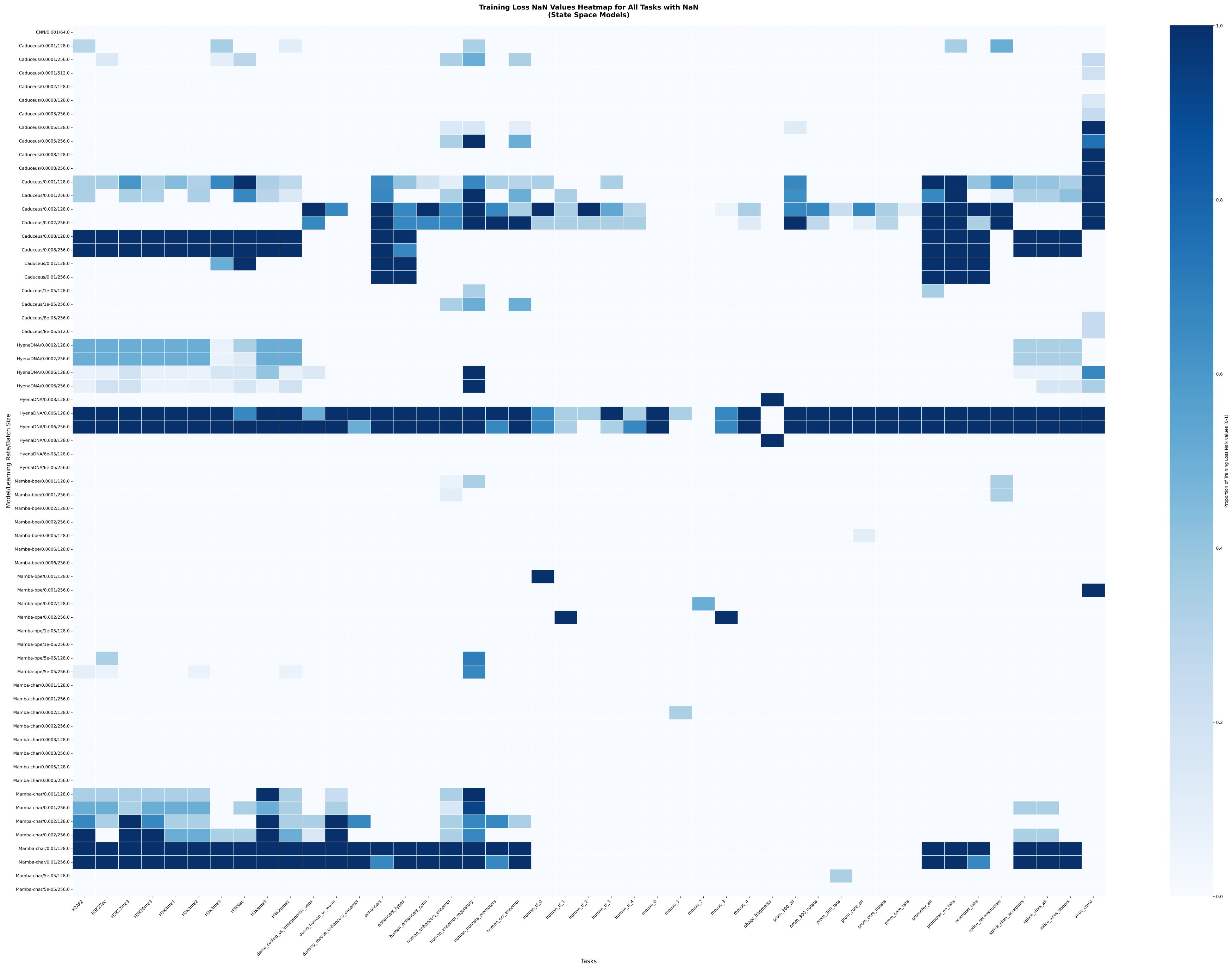


Fig. 2: Training Instability by learning rate and batch size. This heatmap indicates the percentage of instances that resulted in unstable fine-tuning on the genomic benchmark, depending upon the learning rate and batch sizes chosen. Instability is defined here as the number of times that training resulted in a training loss that overflows the precision of the machine and results in a loss of NaN. This typically happens when there is a gradient explosion during training.

#### Complete Benchmarking Results

Accuracy and MCC scores on the Genomic Benchmark can be seen in Tables 7 and 8. The results for the Genome Understanding Evaluation benchmark can be seen in Tables 9 and 10, and the results for the Nucleotide Transformer tasks can be seen in Tables 11 and 12.

**Table 7.** Accuracy Scores on the Genomic Benchmark. The highest score for each dataset is highlighted in bold and underlined.

| | CNN (char) | GPT-2 (bpe) | DNABERT2 (bpe) | Mamba (bpe) | NT (blocked k-mer) | DNABERT (k-mer) | HyenaDNA (char) | Mamba (char) | Caduceus (char) |
|---|---|---|---|---|---|---|---|---|---|
| Mouse Enhancers | 0.715 | 0.679 | 0.716 | 0.775 | 0.693 | 0.630 | **0.795** | 0.785 | 0.788 |
| Coding vs. Intergenic | 0.892 | 0.827 | **0.948** | 0.896 | 0.903 | 0.867 | 0.901 | 0.883 | 0.914 |
| Human vs. Worm | 0.942 | 0.917 | **0.974** | 0.961 | 0.932 | 0.910 | 0.964 | 0.965 | 0.972 |
| Enhancers Cohn | 0.702 | 0.682 | 0.746 | 0.739 | 0.682 | 0.652 | 0.734 | 0.735 | **0.746** |
| Enhancers Ensembl | 0.767 | 0.815 | 0.902 | 0.862 | 0.834 | 0.804 | 0.878 | 0.871 | **0.911** |
| Ensembl Regulatory | 0.866 | 0.883 | 0.771 | 0.892 | **0.937** | 0.771 | 0.861 | 0.733 | 0.869 |
| Non-Tata Promoters | 0.860 | 0.847 | **0.960** | 0.947 | 0.834 | 0.784 | 0.940 | 0.955 | 0.945 |
| OCR Ensembl | 0.695 | 0.634 | 0.812 | 0.772 | 0.686 | 0.631 | 0.802 | 0.816 | **0.817** |
| | *Baseline* | | *Sub-word Tokenization* | | | *Nucleotide Level Tokenization* | | | |

**Table 8.** MCC Scores on the Genomic Benchmark. The highest score for each dataset is highlighted in bold and underlined.

| | CNN (char) | GPT-2 (bpe) | DNABERT2 (bpe) | Mamba (bpe) | NT (blocked k-mer) | DNABERT (k-mer) | HyenaDNA (char) | Mamba (char) | Caduceus (char) |
|---|---|---|---|---|---|---|---|---|---|
| Mouse Enhancers | 0.433 | 0.367 | 0.482 | 0.555 | 0.390 | 0.257 | **0.595** | 0.570 | 0.583 |
| Coding vs. Intergenic | 0.784 | 0.659 | **0.896** | 0.793 | 0.807 | 0.733 | 0.803 | 0.774 | 0.828 |
| Human vs. Worm | 0.884 | 0.836 | **0.949** | 0.923 | 0.865 | 0.820 | 0.929 | 0.930 | 0.943 |
| Enhancers Cohn | 0.409 | 0.365 | 0.493 | 0.478 | 0.368 | 0.305 | 0.469 | 0.473 | **0.493** |
| Enhancers Ensembl | 0.549 | 0.631 | 0.805 | 0.728 | 0.669 | 0.609 | 0.757 | 0.743 | **0.822** |
| Ensembl Regulatory | 0.801 | 0.730 | 0.447 | 0.839 | **0.856** | 0.445 | 0.795 | 0.605 | 0.807 |
| Non-Tata Promoters | 0.721 | 0.703 | **0.920** | 0.895 | 0.673 | 0.586 | 0.880 | 0.911 | 0.891 |
| OCR Ensembl | 0.393 | 0.270 | 0.625 | 0.545 | 0.373 | 0.263 | 0.605 | 0.632 | **0.635** |
| | *Baseline* | | *Sub-word Tokenization* | | | *Nucleotide Level Tokenization* | | | |

**Table 9.** Accuracy Scores on the GUE. The highest score for each dataset is highlighted in bold and underlined.

| | CNN (char) | GPT-2 (bpe) | DNABERT2 (bpe) | Mamba (bpe) | NT (blocked k-mer) | DNABERT (k-mer) | HyenaDNA (char) | Mamba (char) | Caduceus (char) |
|---|---|---|---|---|---|---|---|---|---|
| prom_core_all | 0.808 | 0.815 | 0.816 | 0.798 | 0.771 | **0.853** | 0.813 | 0.808 | 0.828 |
| prom_core_notata | 0.807 | 0.825 | 0.840 | 0.826 | 0.788 | **0.854** | 0.832 | 0.827 | 0.836 |
| prom_core_tata | 0.879 | 0.832 | 0.870 | 0.711 | 0.860 | **0.882** | 0.732 | 0.769 | 0.794 |
| prom_300_all | 0.903 | 0.899 | 0.923 | 0.904 | 0.908 | **0.955** | 0.912 | 0.909 | 0.910 |
| prom_300_notata | 0.936 | 0.944 | **0.966** | 0.955 | 0.927 | 0.964 | 0.959 | 0.958 | 0.964 |
| prom_300_tata | 0.856 | 0.789 | 0.823 | 0.655 | **0.901** | 0.836 | 0.732 | 0.667 | 0.789 |
| human_tfp_0 | 0.784 | 0.821 | 0.829 | 0.815 | 0.777 | 0.830 | **0.830** | 0.830 | 0.826 |
| human_tfp_1 | 0.818 | 0.833 | **0.856** | 0.836 | 0.799 | 0.849 | 0.848 | 0.840 | 0.852 |
| human_tfp_2 | 0.767 | 0.761 | 0.825 | 0.782 | 0.719 | 0.823 | 0.833 | 0.802 | **0.847** |
| human_tfp_3 | 0.678 | 0.686 | **0.790** | 0.736 | 0.649 | 0.773 | 0.757 | 0.739 | 0.766 |
| human_tfp_4 | 0.815 | 0.826 | 0.880 | 0.848 | 0.785 | 0.867 | 0.874 | 0.851 | **0.892** |
| mouse_tfp_0 | 0.645 | 0.662 | 0.797 | 0.752 | 0.669 | 0.726 | 0.760 | 0.731 | **0.803** |
| mouse_tfp_1 | 0.818 | 0.882 | 0.906 | 0.900 | 0.864 | 0.898 | 0.911 | 0.897 | **0.923** |
| mouse_tfp_2 | 0.786 | 0.865 | 0.914 | 0.885 | 0.840 | 0.862 | 0.899 | 0.849 | **0.921** |
| mouse_tfp_3 | 0.688 | 0.760 | 0.880 | 0.792 | 0.712 | 0.826 | 0.866 | 0.741 | **0.911** |
| mouse_tfp_4 | 0.654 | 0.659 | 0.730 | 0.709 | 0.609 | 0.707 | 0.733 | 0.711 | **0.746** |
| virus_covid | 0.227 | 0.601 | 0.470 | **0.691** | 0.555 | 0.456 | 0.663 | 0.625 | 0.672 |
| splice_reconstructed | 0.862 | 0.835 | 0.910 | 0.776 | **0.913** | 0.906 | 0.895 | 0.804 | 0.893 |
| | *Baseline* | | *Sub-word Tokenization* | | | *Nucleotide Level Tokenization* | | | |

**Table 10.** MCC Scores on the GUE. The highest score for each dataset is highlighted in bold and underlined.

| | CNN (char) | GPT-2 (bpe) | DNABERT2 (bpe) | Mamba (bpe) | NT (blocked k-mer) | DNABERT (k-mer) | HyenaDNA (char) | Mamba (char) | Caduceus (char) |
|---|---|---|---|---|---|---|---|---|---|
| prom_core_all | 0.617 | 0.630 | 0.633 | 0.596 | 0.548 | **0.708** | 0.626 | 0.617 | 0.656 |
| prom_core_notata | 0.616 | 0.650 | 0.680 | 0.652 | 0.576 | **0.708** | 0.664 | 0.654 | 0.675 |
| prom_core_tata | 0.761 | 0.670 | 0.744 | 0.424 | 0.722 | **0.764** | 0.465 | 0.540 | 0.595 |
| prom_300_all | 0.807 | 0.798 | 0.847 | 0.810 | 0.816 | **0.909** | 0.826 | 0.818 | 0.821 |
| prom_300_notata | 0.873 | 0.888 | **0.932** | 0.911 | 0.855 | 0.929 | 0.918 | 0.917 | 0.928 |
| prom_300_tata | 0.712 | 0.579 | 0.647 | 0.310 | **0.804** | 0.675 | 0.466 | 0.334 | 0.581 |
| human_tfp_0 | 0.582 | 0.650 | 0.668 | 0.643 | 0.567 | 0.666 | **0.671** | 0.671 | 0.664 |
| human_tfp_1 | 0.643 | 0.674 | **0.718** | 0.683 | 0.605 | 0.703 | 0.706 | 0.691 | 0.716 |
| human_tfp_2 | 0.542 | 0.527 | 0.652 | 0.571 | 0.446 | 0.654 | 0.671 | 0.613 | **0.705** |
| human_tfp_3 | 0.357 | 0.383 | **0.587** | 0.480 | 0.309 | 0.551 | 0.519 | 0.485 | 0.539 |
| human_tfp_4 | 0.639 | 0.656 | 0.764 | 0.697 | 0.573 | 0.739 | 0.749 | 0.706 | **0.786** |
| mouse_tfp_0 | 0.294 | 0.331 | 0.596 | 0.507 | 0.340 | 0.452 | 0.523 | 0.465 | **0.610** |
| mouse_tfp_1 | 0.637 | 0.765 | 0.812 | 0.801 | 0.728 | 0.797 | 0.824 | 0.795 | **0.845** |
| mouse_tfp_2 | 0.574 | 0.731 | 0.829 | 0.772 | 0.682 | 0.726 | 0.801 | 0.701 | **0.845** |
| mouse_tfp_3 | 0.385 | 0.530 | 0.760 | 0.588 | 0.432 | 0.653 | 0.737 | 0.501 | **0.827** |
| mouse_tfp_4 | 0.308 | 0.319 | 0.462 | 0.418 | 0.218 | 0.414 | 0.468 | 0.423 | **0.496** |
| virus_covid | 0.128 | 0.549 | 0.401 | **0.652** | 0.498 | 0.389 | 0.620 | 0.576 | 0.630 |
| splice_reconstructed | 0.766 | 0.718 | 0.846 | 0.614 | **0.852** | 0.840 | 0.820 | 0.664 | 0.816 |
| | *Baseline* | | *Sub-word Tokenization* | | | *Nucleotide Level Tokenization* | | | |

**Table 11.** Accuracy Scores on the Nucleotide Transformer (revised). The highest score for each dataset is highlighted in bold and underlined.

| | CNN (char) | GPT-2 (bpe) | DNABERT2 (bpe) | Mamba (bpe) | NT (blocked k-mer) | DNABERT (k-mer) | HyenaDNA (char) | Mamba (char) | Caduceus (char) |
|---|---|---|---|---|---|---|---|---|---|
| H2AFZ | 0.723 | 0.737 | 0.756 | 0.693 | **0.761** | 0.747 | 0.702 | 0.688 | 0.708 |
| H3K27ac | 0.704 | 0.722 | 0.746 | 0.693 | **0.751** | 0.724 | 0.707 | 0.705 | 0.733 |
| H3K27me3 | 0.746 | 0.771 | 0.799 | 0.752 | **0.801** | 0.774 | 0.764 | 0.757 | 0.763 |
| H3K36me3 | 0.762 | 0.762 | 0.811 | 0.738 | **0.828** | 0.778 | 0.746 | 0.737 | 0.760 |
| H3K4me1 | 0.690 | 0.711 | 0.749 | 0.693 | **0.752** | 0.727 | 0.695 | 0.692 | 0.706 |
| H3K4me2 | 0.743 | 0.758 | **0.782** | 0.729 | 0.777 | 0.763 | 0.732 | 0.724 | 0.736 |
| H3K4me3 | 0.789 | 0.790 | 0.816 | 0.800 | **0.822** | 0.808 | 0.799 | 0.802 | 0.811 |
| H3K9ac | 0.751 | 0.746 | 0.776 | 0.737 | **0.778** | 0.750 | 0.751 | 0.749 | 0.755 |
| H3K9me3 | 0.666 | 0.691 | 0.746 | 0.643 | **0.749** | 0.714 | 0.681 | 0.668 | 0.697 |
| H4K20me1 | 0.776 | 0.791 | 0.821 | 0.766 | **0.830** | 0.800 | 0.782 | 0.775 | 0.788 |
| promoter_all | 0.867 | 0.852 | 0.878 | 0.848 | **0.892** | 0.864 | 0.854 | 0.854 | 0.859 |
| promoter_no_tata | 0.864 | 0.869 | 0.880 | 0.863 | **0.896** | 0.869 | 0.867 | 0.866 | 0.868 |
| promoter_tata | **0.943** | 0.890 | 0.928 | 0.808 | 0.942 | 0.917 | 0.834 | 0.836 | 0.919 |
| enhancers | 0.745 | 0.731 | 0.755 | 0.736 | **0.797** | 0.761 | 0.740 | 0.737 | 0.745 |
| enhancers_types | 0.705 | 0.698 | 0.727 | 0.700 | **0.771** | 0.728 | 0.703 | 0.709 | 0.709 |
| splice_site_prediction | 0.951 | 0.831 | 0.903 | 0.803 | **0.978** | 0.974 | 0.939 | 0.953 | 0.941 |
| splice_sites_acceptors | 0.964 | 0.868 | 0.904 | 0.802 | **0.985** | 0.979 | 0.944 | 0.951 | 0.937 |
| splice_sites_donors | 0.976 | 0.874 | 0.915 | 0.797 | **0.986** | 0.983 | 0.960 | 0.960 | 0.940 |
| | *Baseline* | | *Sub-word Tokenization* | | | *Nucleotide Level Tokenization* | | | |

#### Genome Benchmark

The Genomic Benchmark was introduced by Gresova et al [5] in 2023. This benchmark includes identification of regulatory elements in human DNA, discriminating between coding and non-coding DNA, identification of regulatory elements in mouse and drosophilia DNA and discriminating between Human and Worm DNA.

**Human Enhancers Cohn** This dataset was adapted from Cohn et al. 2018 [2]. This consists of simulated sequences, in vivo binding data for individual transcription factors, and genome-wide chromatin maps of active enhancers.

**Human Enhancers Ensembl** This dataset includes human enhancers from The FANTOM5 project by Andersson et al. 2014 [1]. The human genome GRCh38 was used to randomly generate negative sequences that match the lengths of the positive sequences without overlap.

**Human Ensemble Regulatory** This dataset was retrieved from Howe et al. 2021 [6] which contains human enhancer, promoter, and open chromatin region sequences from Zerbino et al 2015 [13].

**Human Non-tata Promoters** This dataset was curated by Umarov et al. [12] and consists of promoters from human, mouse, plant (Arabidopsis), and the bacteria Escherichia coli and Bacillus subtilis. E. coli promoters were extracted from RegulonDB and Bacillus subtilus promoters were taken from DBTBS [7]. Both positive and negative examples were 81 nucleotides in length. The negative samples

**Table 12.** MCC Scores on the Nucleotide Transformer (revised). The highest score for each dataset is highlighted in bold and underlined.

| | CNN (char) | GPT-2 (bpe) | DNABERT2 (bpe) | Mamba (bpe) | NT (blocked k-mer) | DNABERT (k-mer) | HyenaDNA (char) | Mamba (char) | Caduceus (char) |
|---|---|---|---|---|---|---|---|---|---|
| H2AFZ | 0.455 | 0.483 | 0.525 | 0.405 | **0.526** | 0.504 | 0.413 | 0.392 | 0.432 |
| H3K27ac | 0.409 | 0.445 | 0.500 | 0.392 | **0.504** | 0.455 | 0.421 | 0.418 | 0.467 |
| H3K27me3 | 0.509 | 0.555 | 0.609 | 0.530 | **0.619** | 0.568 | 0.542 | 0.535 | 0.551 |
| H3K36me3 | 0.540 | 0.539 | 0.627 | 0.489 | **0.662** | 0.572 | 0.504 | 0.486 | 0.531 |
| H3K4me1 | 0.406 | 0.431 | 0.506 | 0.406 | **0.513** | 0.467 | 0.403 | 0.405 | 0.429 |
| H3K4me2 | 0.487 | 0.534 | **0.575** | 0.470 | 0.562 | 0.536 | 0.477 | 0.456 | 0.485 |
| H3K4me3 | 0.581 | 0.581 | 0.634 | 0.601 | **0.644** | 0.616 | 0.600 | 0.606 | 0.623 |
| H3K9ac | 0.503 | 0.495 | 0.555 | 0.477 | **0.556** | 0.504 | 0.505 | 0.502 | 0.512 |
| H3K9me3 | 0.344 | 0.388 | 0.495 | 0.290 | **0.499** | 0.433 | 0.370 | 0.340 | 0.395 |
| H4K20me1 | 0.560 | 0.588 | 0.657 | 0.544 | **0.674** | 0.609 | 0.574 | 0.560 | 0.589 |
| promoter_all | 0.735 | 0.704 | 0.758 | 0.698 | **0.785** | 0.728 | 0.708 | 0.711 | 0.718 |
| promoter_no_tata | 0.731 | 0.738 | 0.761 | 0.727 | **0.793** | 0.740 | 0.735 | 0.733 | 0.738 |
| promoter_tata | **0.888** | 0.784 | 0.858 | 0.621 | 0.885 | 0.834 | 0.671 | 0.676 | 0.838 |
| enhancers | 0.501 | 0.475 | 0.523 | 0.487 | **0.599** | 0.539 | 0.492 | 0.489 | 0.505 |
| enhancers_types | 0.442 | 0.447 | 0.493 | 0.449 | **0.572** | 0.495 | 0.447 | 0.462 | 0.461 |
| splice_site_prediction | 0.927 | 0.748 | 0.854 | 0.705 | **0.967** | 0.961 | 0.909 | 0.930 | 0.911 |
| splice_sites_acceptors | 0.928 | 0.736 | 0.808 | 0.607 | **0.971** | 0.957 | 0.887 | 0.902 | 0.873 |
| splice_sites_donors | 0.953 | 0.750 | 0.829 | 0.598 | **0.973** | 0.967 | 0.919 | 0.919 | 0.880 |
| | *Baseline* | | *Sub-word Tokenization* | | | *Nucleotide Level Tokenization* | | | |

were generated by randomly selecting fragments of protein-coding genes and then taking the reverse complement of the strand. Eukaryotic promoters were extracted from EPD [4]. Negative examples were randomly generated segments of length 251 from the gene located after the first exon.

**Human OCR Ensembl** This dataset was adapted from Howe et a. 2021 [6] and Zerbino et al. 2015 [13]. The positive examples in this dataset are human open chromatin regions that were seen in experiments using DNase-seq but are not included in any other categories like enhancer, promoter, gene, TSS, or CTCF [13]. Using the human genome GRCh38, negative examples were randomly generated to match the lengths of the positive sequences without overlap.

**Human or Worm (demo)** This is a demo dataset compiled by Gresova et al. [5]. It consists of randomly selected 200 base pair long human and C. elegans genomic sequences with a 75:25 train-test split.

**Coding or Intergenomic (demo)** This is a dataset compiled of randomly selected 200 base pair intervals from intergenomic regions or protein coding transcipts of the human genome with a 75:25 train-test split. The datasets are from the simecek/PseudoDNA_Generator GitHub repo [11] and were used in the ECCB deep learning workshop.

**Mouse Enhancers Ensembl (dummy)** The dataset was curated using data from Ensembl release 100. Positive sequences consist of mouse enhancers sourced from the VISTA Enhancer Browser, while negative sequences were randomly generated from the mouse genome GRCm38 to match the lengths of the positive sequences without overlapping.

#### Nucleotide Transformer Tasks (revised)

A set of genomic classification tasks was introduced by Dalle-Torre et al [3]. This version of the benchmark supercedes the original version. It contains ten binary classification tasks of epigenetic mark prediction from a human dataset, a dataset of regulatory elements from multiple species and a splice site detection dataset from human DNA. In all there are 18 fine-tuning tasks with either two or three labeled classes.

**Promoter Detection (Human)** This dataset was constructed by the authors by downloading all human promoters from the Eukaryotic Promoter Database [9], selecting the region that spans 49 nt upstream and 10 nt downstream from the transcription start sites. They then selected 300-nt regions that contained promoters as positive examples and all remaining 300-nt regions as negative examples. These were then split to create three different tasks: the presence of any promoter element (promoter all), a promoter with a tata box or a promoter without a tata box.

**Enhancers (Human)** Human enhancer elements were downloaded from the ENCODE's SCREEN database [8] and and split into tissue-specific or tissue-invariant. Regions 400 nt in length containing enhancers were selected as positive and not overlapping enhancers were labeled negative. The enhancer task is a binary classification task, with the model identifying a segment as containing or not containing an enhancer. The enhancer types dataset is a multi-label prediction task with the three labels: tissue-specific enhancer, tissue-invariant enhancer or none.

**Splice Site Prediction (Human)** Human annotated splice sites were downloaded from GENCODE and filtered to exclude level-3 transcripts to obtain the highest quality training dataset. Splice site annotations were extracted and then 600 nt genomic regions were selected with the positive set containing a splice donor or acceptor site in the center of the window and all remaining 600 nt segments that were not overlapping a splice site were labeled as negative. Three different tasks were created from this dataset, a binary classification task for donors, acceptors and a multi-label prediction task with the labels being acceptor, donor or none.

**Epigenetic Marks Prediction (Human)** This data was curated by the authors by downloading histone ChIP-seq data for ten histone marks in the K562 human cell line from ENCODE [8] A 1-kb genomic sequence containing peaks were selected for each positive example and all 1-kb sequences that were not overlapping any peaks were selected as negative examples.

**Dataset splits and subsampling** Each dataset was balanced by subsampling the negative examles to the same number as positive examples. To reduce the overal size of the datasets, the datasets were further randomly sampled to create a training dataset of size 30000 and a validation and test dataset of size 3000.

#### Genome Understanding Evaluation

The authors of DNABERT-2 introduced a new Genomic Understanding Evaluation (GUE) Benchmark in 2023 [15]. They felt that previously published genomic benchmarks were too easy for gLM's and so they rebalanced the datasets that they constructed to have an accuracy of between 0.3 and 0.9, to provide a more challenging benchmark for gLMs. The epigenetic marks prediction tasks are based on the same dataset as the Nucleotide Transformer Tasks version 1, and so we do not provide repeated results for these datasets, instead providing results for the epigenetic marks prediction tasks in NTTv2 because these datasets have been curated and balanced and are more robust than the originals. They also introduce tasks of human transcription factor prediction, mouse transcription factor prediction, and a challenging covid variant classification task with nine classes.

**Promoter Detection (Human)** The data for these three datasets came from the Eukaryotic Promoter Dataset (EPDnew), the same primary source as the Nucleotide Transformer benchmark [3].

**Core Promoter Detection (Human)** Similar to the Promoter Detection dataset, but the window of nucleotides included are -34 and +35 from the TSS, which should make this a more challenging task than proximal promoter prediction.

**Splice Site Prediction (Human)** The data for this task was originally compiled by Wang et al. 2019. The dataset consists of positive and negative examples of donor, acceptor and non-splice sites that were extracted from Ensemble GRCh38. This is the same source as the original splice_sites_all task from Dalla-Torre et al. 2023 [3], but the dataset was reconstructed by Zhou et al. 2024 with more adversarial non-cannonical splice site examples to make it more challenging.

**Transcription Factor Binding Site Prediction (Human)** This dataset was constructed by downloading 690 ENCODE ChIP-seq experiments from 2012 and extracted a 101 bp region around the center of each peak for the true positive class and extracted sequences that were not overlapping of the same length for the negative class. Five datasets were randomly constructed from this set and they were curated heuristically so that the accuracy range would have an F1 score between 0.5 and 0.95.

**Transcription Factor Binding Site Prediction (Mouse)** Similar to the TFBS prediction for human DNA, but mouse ENCODE ChIP-seq data came from Stamatoyannopoulos et al. 2012, and negative examples were created using dinucleotide shuffling while preserving relative frequencies.

**Covid Variant Prediction (Virus)** This dataset was compiled by Zhou et al. [15]. They downloaded viral genomes from EpiCov database from GISAID. They extract 1000 bp segments from each genome and classify nine variants of the virus, Alpha, Beta, Delta, Eta, Gamma, Iota, Kappa, Lambda and Zeta.

**Enhancer promoter interaction (Human)** This dataset was compiled by Zhou et al. [15] and discerns the interactions between promoters and enhancers.

#### References

1. Robin Andersson, Claudia Gebhard, Irene Miguel-Escalada, Ilka Hoof, Jette Bornholdt, Mette Boyd, Yun Chen, Xiaobei Zhao, Christian Schmidl, Takahiro Suzuki, Evgenia Ntini, Erik Arner, Eivind Valen, Kang Li, Lucia Schwarzfischer, Dagmar Glatz, Johanna Raithel, Berit Lilje, Nicolas Rapin, Frederik Otzen Bagger, Mette Jørgensen, Peter Refsing Andersen, Nicolas Bertin, Owen Rackham, A. Maxwell Burroughs, J. Kenneth Baillie, Yuri Ishizu, Yuri Shimizu, Erina Furuhata, Shiori Maeda, Yutaka Negishi, Christopher J. Mungall, Terrence F. Meehan, Timo Lassmann, Masayoshi Itoh, Hideya Kawaji, Naoto Kondo, Jun Kawai, Andreas Lennartsson, Carsten O. Daub, Peter Heutink, David A. Hume, Torben Heick Jensen, Harukazu Suzuki, Yoshihide Hayashizaki, Ferenc Müller, Alistair R. R. Forrest, Piero Carninci, Michael Rehli, and Albin Sandelin. An atlas of active enhancers across human cell types and tissues. *Nature*, 507(7493):455–461, March 2014.
2. Dikla Cohn, Or Zuk, and Tommy Kaplan. Enhancer identification using transfer and adversarial deep learning of DNA sequences, February 2018. Pages: 264200 Section: New Results.
3. Hugo Dalla-Torre, Liam Gonzalez, Javier Mendoza-Revilla, Nicolas Lopez Carranza, Adam Henryk Grzywaczewski, Francesco Oteri, Christian Dallago, Evan Trop, Bernardo P. de Almeida, Hassan Sirelkhatim, Guillaume Richard, Marcin Skwark, Karim Beguir, Marie Lopez, and Thomas Pierrot. The Nucleotide Transformer: building and evaluating robust foundation models for human genomics, October 2024. bioRxiv: 10.1101/2023.01.11.523679.
4. René Dreos, Giovanna Ambrosini, Rouayda Cavin Périer, and Philipp Bucher. EPD and EPDnew, high-quality promoter resources in the next-generation sequencing era. *Nucleic Acids Research*, 41(Database issue):D157–164, January 2013.
5. Katarína Grešová, Vlastimil Martinek, David Čechák, Petr Šimeček, and Panagiotis Alexiou. Genomic benchmarks: a collection of datasets for genomic sequence classification. *BMC Genomic Data*, 24(1):25, May 2023.
6. Kevin L Howe, Premanand Achuthan, James Allen, Jamie Allen, Jorge Alvarez-Jarreta, M Ridwan Amode, Irina M Armean, Andrey G Azov, Ruth Bennett, Jyothish Bhai, Konstantinos Billis, Sanjay Boddu, Mehrnaz Charkhchi, Carla Cummins, Luca Da Rin Fioretto, Claire Davidson, Kamalkumar Dodiya, Bilal El Houdaigui, Reham Fatima, Astrid Gall, Carlos Garcia Giron, Tiago Grego, Cristina Guijarro-Clarke, Leanne Haggerty, Anmol Hemrom, Thibaut Hourlier, Osagie G Izuogu, Thomas Juettemann, Vinay

Kaikala, Mike Kay, Ilias Lavidas, Tuan Le, Diana Lemos, Jose Gonzalez Martinez, José Carlos Marugán, Thomas Maurel, Aoife C McMahon, Shamika Mohanan, Benjamin Moore, Matthieu Muffato, Denye N Oheh, Dimitrios Paraschas, Anne Parker, Andrew Parton, Irina Prosovetskaia, Manoj P Sakthivel, Ahamed I Abdul Salam, Bianca M Schmitt, Helen Schuilenburg, Dan Sheppard, Emily Steed, Michal Szpak, Marek Szuba, Kieron Taylor, Anja Thormann, Glen Threadgold, Brandon Walts, Andrea Winterbottom, Marc Chakiachvili, Ameya Chaubal, Nishadi De Silva, Bethany Flint, Adam Frankish, Sarah E Hunt, Garth R IIsley, Nick Langridge, Jane E Loveland, Fergal J Martin, Jonathan M Mudge, Joanella Morales, Emily Perry, Magali Ruffier, John Tate, David Thybert, Stephen J Trevanion, Fiona Cunningham, Andrew D Yates, Daniel R Zerbino, and Paul Flicek. Ensembl 2021. *Nucleic Acids Research*, 49(D1):D884–D891, January 2021.

7. Takahiro Ishii, Ken-ichi Yoshida, Goro Terai, Yasutaro Fujita, and Kenta Nakai. DBTBS: a database of Bacillus subtilis promoters and transcription factors. *Nucleic Acids Research*, 29(1):278–280, January 2001.
8. Y. Luo, B.C. Hitz, I. Gabdank, J.A. Hilton, M.S. Kagda, B. Lam, Z. Myers, P. Sud, J. Jou, K. Lin, U.K. Baymuradov, K. Graham, C. Litton, S.R. Miyasato, J.S. Strattan, O. Jolanki, J.W. Lee, F.Y. Tanaka, P. Adenekan, E. O'Neill, and J.M. Cherry. New developments on the Encyclopedia of DNA Elements (ENCODE) data portal. *Nucleic Acids Research*, 48(D1):D882–D889, January 2020.
9. Rouaïda Cavin Périer, Viviane Praz, Thomas Junier, Claude Bonnard, and Philipp Bucher. The Eukaryotic Promoter Database (EPD). *Nucleic Acids Research*, 28(1):302–303, January 2000.
10. Yair Schiff, Chia-Hsiang Kao, Aaron Gokaslan, Tri Dao, Albert Gu, and Volodymyr Kuleshov. Caduceus: Bi-directional equivariant long-range DNA sequence modeling. *arXiv preprint arXiv:2403.03234*, 2024.
11. Petr Simecek. PseudoDNA Generator, July 2020. https://github.com/simecek/ PseudoDNA_Generator.
12. Ramzan Kh Umarov and Victor V. Solovyev. Recognition of prokaryotic and eukaryotic promoters using convolutional deep learning neural networks. *PLoS One*, 12(2):e0171410, February 2017.
13. Daniel R. Zerbino, Steven P. Wilder, Nathan Johnson, Thomas Juettemann, and Paul R. Flicek. The ensembl regulatory build. *Genome Biology*, 16(1):56, March 2015.
14. Jingjing Zhai, Aaron Gokaslan, Yair Schiff, Ana Berthel, Zong-Yan Liu, Wei-Yun Lai, Zachary R. Miller, Armin Scheben, Michelle C. Stitzer, M. Cinta Romay, Edward S. Buckler, and Volodymyr Kuleshov. Cross-species modeling of plant genomes at single nucleotide resolution using a pre-trained DNA language model. *bioRxiv: The Preprint Server for Biology*, page 2024.06.04.596709, August 2024.
15. Zhihan Zhou, Yanrong Ji, Weijian Li, Pratik Dutta, Ramana Davuluri, and Han Liu. DNABERT-2: efficient foundation model and benchmark for multi-species genome, June 2023. arXiv: 2306.15006.
